# From Discovery Proteomics to Process-Informed Monitoring in Biomanufacturing Chassis Development

**DOI:** 10.64898/2025.12.18.695173

**Authors:** Matthew R. Russell, Philip J. Brownridge, Joseph Windo, Nigel S. Scrutton, Claire E. Eyers, Perdita Barran

## Abstract

Gaining control of existing biomanufacturing chassis organisms, such as *Escherichia coli K12*, and novel isolates, such as the salt tolerant *Halomonas bluephagenesis sp TD01* studied here may be facilitated by the investigation and monitoring of their metabolic and regulatory processes, particularly through proteomics. Here we consider the performance of a range of typically available proteomics platforms across a range of price points to map chassis organisms’ metabolic pathways. A set of model bacterial samples was prepared from *E. coli* and *H. bluephagenesis sp. TD01* in 1:1, 1:2 and 2:1 ratios and analyzed using five LC-MS systems. Of the 8,222 proteins identified across all samples analysed (4,401 proteins from *E. coli*; 3,821 from *Halomonas sp. TD01*), the TimsTOF and Exploris were able to achieve extensive proteome coverage quantifying 5.5k and 5k proteins respectively, with the ZenoTOF, Waters MRT and the legacy Waters Vion respectively quantifying 3.5k, 1.3k, and ∼850 proteins at 1% FDR. Proteins comprising core metabolic pathways critical to biomanufacturing in these chassis’ organisms can be quantified with all instruments. We characterize metabolic adaptation in *H. bluephagenesis* by showing that replacement of glucose with a carboxylic acid feed stock directs carbon flux towards potential butane precursors as well as how the acquired data permits monitoring of the cobalamin (vitamin B_12_) production pathway.

**TOC Graphic:** 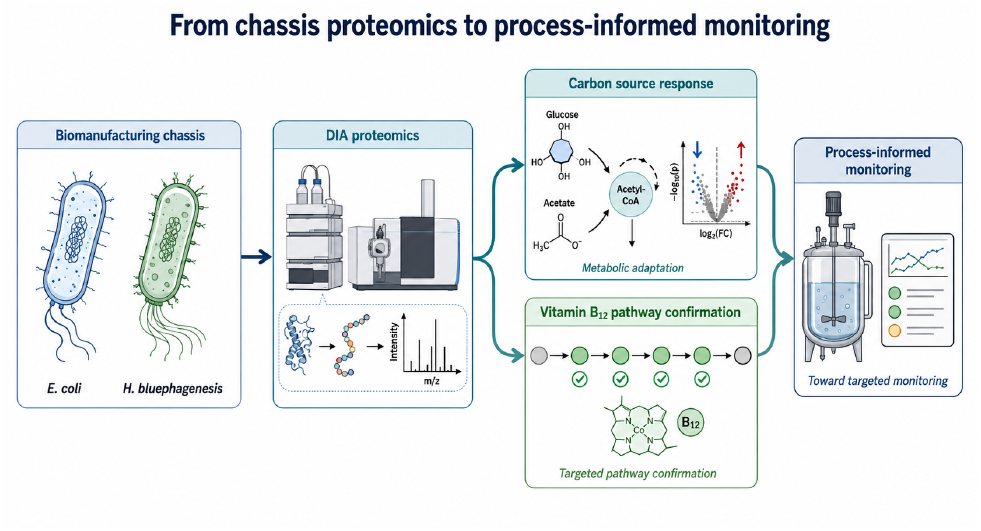

## Background

The transition toward a sustainable bioeconomy demands efficient biomanufacturing platforms capable of producing low-value, high-volume bulk commodities such as hydrocarbons^1^. Biomanufacturing which puts living organisms to work producing chemical products at industrial scale^2,3^ is an economically important industrial sector with an estimated global value of $19.08 billion in 2024 ^4^. While *Escherichia coli* (*E. coli*) remains the most extensively characterized microbial chassis, its metabolic and physiological limitations preclude its use in certain industrial contexts, necessitating the development of alternative host organisms. Compared to *E. coli* such candidates lack the extensive biochemical and microbiological research base accumulated over decades and this knowledge gap must be addressed before they can be rationally engineered for production. *Halomonas bluephagenesis* (*H. bluephagenesis*) exemplifies both the promise and the challenge of such emerging chassis organisms^5–7^ As a relatively recently isolated halophilic species, *H. bluephagenesis* has attracted considerable interest in biomanufacturing applications owing to its capacity to grow in high-salt, high-pH media — conditions inhospitable to most terrestrial microbial contaminants^8^. This natural tolerance raises the prospect of non-sterile fermentation, potentially eliminating costly sterilization steps in industrial-scale production^9^. Nevertheless, the research base for *H. bluephagenesis* remains less developed than that of *E. coli*, and here we employ it as a case study for any novel chassis that the biotechnology sector may seek to adopt.

Systems biology approaches, and proteomics in particular, offer a route to rapidly characterize the physiology of such organisms under biomanufacturing-relevant conditions. Proteomics provides a systems level overview of the protein machinery of biological systems yet the application of proteomics to biomanufacturing remains under explored compared to medicine^10^ or indeed to microbiology in general^11,12^ although it has shown utility when applied to biopharmaceutical production^13,14^. Quantitative proteomics enables the mapping of protein expression onto metabolic pathways curated in databases such as KEGG, providing insight into how organisms respond to growth conditions, heterologous pathway expression, and other operational perturbations. When integrated with genome-scale metabolic models, such data could further inform rational strategies for metabolic engineering.

Data independent acquisition (DIA) proteomics is particularly attractive for biomanufacturing applications because it provides reproducible quantitative measurements across large sample series and over long time periods. In DIA workflows, peptide ions within sequential precursor windows are systematically subjected to collisions that cause the production of diagnostic fragments during chromatographic elution. This enables consistent peptide detection and quantification between experiments. DIA was selected rather than data dependent acquisition (DDA) because the long-term objective of this work is the development of reproducible targeted workflows suitable for process monitoring and industrial deployment. Compared with DDA approaches, DIA generally provides improved quantitative reproducibility, reduced missing values across large sample cohorts, and more consistent peptide detection between runs. These characteristics are particularly important in biomanufacturing settings where process deviations may need to be detected over long operational timeframes or across multiple sites. Different vendors implement DIA using distinct label free acquisition strategies, including ion mobility-assisted methods such as diaPASEF and HDMSe, conventional SWATH approaches, and Orbitrap-based DIA methods^15–17^. In this study, we evaluated the performance of mass spectrometry platforms for proteomic analysis of microbial chassis organisms. Our intention was not to establish the theoretical maximum performance achievable on each platform through extensive expert optimization, but rather to compare representative workflows likely to be implemented by users adopting DIA proteomics in biomanufacturing environments. Accordingly, we used either manufacturer-recommended or standardized acquisition strategies for each platform, reflecting realistic deployment conditions and minimizing operator-dependent optimization bias.

To provide a well-characterised reference for assessing representative workflow performance, we employed *E. coli* as a well-characterized reference proteome, alongside *H. bluephagenesis* as a representative emerging chassis^18^. By generating comprehensive proteomic datasets for both organisms, we were able to assess not only instrument-level coverage and sensitivity, but also the applicability of these platforms to addressing genuine biotechnological questions — specifically, the adaptive response of H. bluephagenesis to changes in carbon feedstock and the verification of native vitamin B_12_ pathway expression.

To assess the various instruments’ performance, under standard operating conditions we first prepared mixtures of peptides from E. coli and H. bluephagenesis in ratios of 1:1, 1:2 and 2:1 in a similar approach to that adopted in multi-site comparison studies^16,17^. These ratios were selected because the 2:1 and 1:2 samples should produce a set 2-fold difference in “expression” which is usually considered the smallest biologically significant degree of differential expression and therefore represents the most difficult real-world challenge to detect differential expression. Furthermore, larger degrees of differential expression, ten-fold for example, could push the limit of the dynamic range of instruments.

Data-independent acquisition (DIA) liquid chromatography mass spectrometry data was acquired for aliquots of each standard, on each of the instruments described above. The numbers of proteins identified, quantified, and the data’s sensitivity to differential expression are reported for each instrument. Our intention was not to identify a “best” instrument, nor to finesse acquisition conditions, but rather to show the value that can be obtained irrespective of platform, as relevant to advance biomanufacturing, and to provide model datasets and robust methodology for such applications. As a worked example, we showcase how the Sciex ZenoTOF, ranking middle in terms of both price and performance, can support bioengineering development in *H. bluephagenesis* for biofuel production. Specifically, we were able to demonstrate that replacement of glucose with a carboxylic acid feed stock - a low-cost byproduct of biodiesel production – directs carbon flux towards potential butane precursors, characterizing the metabolic adaptation of *H. bluephagenesis*. We also demonstrate that such data may be used to confirm expression of coenzyme across a pathway and here apply it to the production of cobalamin (vitamin B_12_).

This integrated experimental design thus serves a dual purpose: it provides a systematic comparison of mass spectrometry instrumentation for microbial proteomics, while simultaneously demonstrating how such platforms can accelerate the characterization and engineering of emerging industrial chassis organisms. We envisage the data sets produced will facilitate industrial adoption of such techniques on other sites by providing reference data alongside processing scripts and example results. The limited adoption of proteomics by the biomanufacturing industry can be attributed to startup costs, lack of skills, and the risk that projects fail to positively impact the bottom line. Costs include capital investment in proteomics facilities and equipment and staffing for skilled operators. Legitimate doubts about the payoff remain given the absence of cases studies complete with reference data, methodology and workflows that demonstrate the benefits of application of this technology to increase product yield and/or lower production costs^19^.

## Results

### Protein Identification and Quantification across Five Platforms

A model sample set comprising ratio mixtures of 1:2, 1:1 and 2:1 by total peptide mass, of *E. coli* and *Halomonas bluephagenesis* sp TD01 whole cell lysates was analyzed on current high end mass spectrometry instrumentation: Bruker timsTOF HT; Thermo Exploris 480; and Sciex ZenoTOF; a modern bench top instrument pitched for low cost routine measurement: Waters Xevo MRT; and a legacy instrument developed for MPK and CRO work Waters Vion. Quantitative data was acquired in DIA mode for all instruments because that technique is the most suitable for longitudinal monitoring of bioreactor conditions and subsequent industrial production. DIA facilitates comparison of quantitative results accumulated over time^15,16^ and potentially at differing sites^17^. Three of the instruments (timsTOF, Exploris and MRT) were equipped with Evosep LC systems which have a limited set of pre-programmed LC methods, enabling identical LC methods to be applied without the need for optimization for each instrument. The other two instruments (Vion and zenoTOF) were equipped with Waters’ Waters Acquity M-Class LC systems which could not be set up identically to the Evosep systems, we therefore worked to optimize our data acquisition methods for these instruments to compare them at optimal performance, using pre-programmed methods on the Evosep. Data from all instruments, apart from the Vion - for which it was not possible - were processed through the same data processing pipeline of DIANN and MSstats using identical libraries. Settings were kept as close as possible and only modified where particular features of the instrument - such as mass accuracy - meant the standard parameter would unfairly handicap the instrument’s apparent performance. This study should be interpreted as a representative workflow comparison rather than a comprehensive instrument benchmark. We did not optimize each platform for our samples or include every current proteomics instrument, instead, we asked what information could be obtained using practical, in-house, DIA workflows across instruments spanning different levels of performance, price and accessibility. Biological examples are included to demonstrate how such data can inform chassis development and targeted pathway monitoring, rather than to provide a complete mechanistic analysis of H. bluephagenesis metabolism.

The protein identification numbers for each of the conditions are tabulated in Table 1, percent sequence coverage for each protein, for each instrument is found in supplementary data set 1. Bacteria, of the kind used for biomanufacturing, typically have genomes coding for between 500 and 5,000 proteins. The fasta protein sequences used here contain 4,401 and 3,821 proteins for *E. coli* and *H. bluephagenesis*respectively yielding a total of 8,222 proteins potentially detectable in the combined samples. Peptides from all of these 8,222 proteins were included in the spectral library generated by DIANN. The full complement of proteins coded for by a bacterial genome will not be expressed; for example proteins required only for spore formation are not likely to be expressed in fully active cells in liquid media. The quantification of 4,726 and 5,554 proteins respectively using either the Exploris 480 or the timsTOF HT instruments represents near complete coverage of the expressed genome of these two species under the sampling conditions. We suggest that the slightly higher number of counts from the timsTOF is due to the additional separation of peptides by ion mobility. Fewer proteins (3,402) were quantifiable from the ZenoTOF data, albeit acquired with a shorter LC gradient. While it is possible that higher coverage could have been achieved with this instrument using a longer LC gradient, we elected to use manufacturer recommended methods than exhaustively optimize conditions for all five platforms. This decision is in line with our study design, which is to demonstrate what could be done in a biomanufacturing setting using routine recommended methods, on instruments that may have other primary purposes. The remaining two instruments permitted quantification of 969 and 862 proteins for the Xevo MRT and the Vion respectively, equating to 17% and 15% of the numbers identified using the timsTOF.

**Table 1:** Summary of workflow-level performance characteristics across the five LC-MS platforms with indicative price points stated. The comparison integrates protein and peptide identification depth, reproducibility, quantitative performance, biological/pathway coverage and practical deployment considerations. Values are derived from the ratio-mixture workflow unless otherwise stated. The table is intended to summarize representative deployable workflow performance rather than define absolute instrument capability.

| Platform and (price point) | Acquisition / workflow features | Proteins quantified at 1% FDR | Peptides quantified | Reproducibility / repeatability | Quantitative performance | Biological / pathway coverage | Practical interpretation |
| --- | --- | --- | --- | --- | --- | --- | --- |
| Bruker timsTOF HT (High) | diaPASEF; ion mobility-assisted DIA; high sequencing speed; gas-phase separation reduces spectral complexity | 5554 | 71738 | High repeatability group; ~75 85% of proteins identified across replicate injections | Strong fold-change performance; high sensitivity for differentially abundant proteins; contributes to median observed fold-change of 1.76 across platforms for nominal 2-fold mixtures | Highest KEGG pathway coverage; carbon metabolism coverage ~91%; ABC transporter coverage >80% | Best suited for deep discovery, strain characterisation and early process-development studies where maximal proteome depth is required |
| Thermo Exploris 480 (High) | Orbitrap DIA; high resolution; narrow precursor isolation windows; Evosep workflow | 4726 | 53836 | High repeatability group; ~75 85% of proteins identified across replicate injections | Strongest statistical confidence for detecting specified protein ratios; high quantitative accuracy | Extensive KEGG pathway coverage; strong PaxDb agreement; broad coverage of high- and lower-abundance proteins | High-performance discovery platform suitable for deep pathway mapping and generation of candidate markers for later targeted monitoring |
| Sciex ZenoTOF 7600 (Mid) | SWATH-MS with Zeno trap; mid-tier platform; shorter LC gradient than some high-end workflows | 3402 | 28163 | High repeatability group; ~75 85% of proteins identified across replicate injections | Good quantitative performance despite lower depth than timsTOF/Exploris; suitable for biologically interpretable differential analysis | Sufficient depth for pathway-level interpretation; quantified 2,818 H. bluephagenesis proteins in the glucose/acetate exemplar | Selected for biological exemplar because it represents an accessible mid-tier workflow capable of process-relevant biological insight. |

Table 1: Summary of workflow-level performance characteristics across the five LC-MS platforms with indicative price points stated cont.
| Platform and (price point) | Acquisition / workflow features | Proteins quantified at 1% FDR | Peptides quantified | Reproducibility / repeatability | Quantitative performance | Biological / pathway coverage | Practical interpretation |
| --- | --- | --- | --- | --- | --- | --- | --- |
| Waters Xevo MRT (Mid) | Benchtop multi-reflectron TOF; MSE-style acquisition; early-access proteomics workflow; Evosep LC | 969 | 5248 | Lower repeatability group; ~52% protein-level repeatability | Ratio compression observed, likely reflecting background noise and less experience with optimisation for proteomic applications | Partial but useful coverage of core pathways; lower abundance proteins less consistently quantified | Represents lower-cost/benchtop deployable instrumentation; useful for selected pathway monitoring but less suited to broad discovery in current configuration |
| Waters Vion (Mid and Legacy) | Legacy IMS-QTOF HDMSE ion mobility enabled workflow; alternative Skyline-based processing route required | 862 | 3376 | Lower repeatability group; ~52% protein-level repeatability | Lowest depth; quantitative reproducibility sufficient for a subset of higher-abundance proteins | Despite limited depth, captured ~43% of carbon metabolism proteins, illustrating utility for selected biological questions | Demonstrates that even legacy/workhorse platforms can report biologically informative subsets of the proteome, but are less suited to comprehensive discovery |

Table 1 summarizes the main performance characteristics of each platform across multiple dimensions: protein and peptide identification depth, replicate consistency, quantitative performance, abundance-range behavior, pathway coverage and practical deployment. Reproducibility of protein and peptide level quantification for each of the instruments across replicate injections is presented in Figure 1 A-C. At the protein level Figure 1 A, replication was more consistent across the timsTOF HT, Exploris 480 and ZenoTOF platforms, with 75-85% of proteins identified in at least one of the nine replicate injections. These numbers decrease to ∼52% for the percentage repeatably identified on the Vion and the MRT. As expected, data from the lowest sensitivity instrument (Vion) provided the fewest quantified proteins. For the MRT we recognize that this new platform has had less optimization for proteomics applications and expect higher numbers and better reproducibility may be possible. Nevertheless, coverage even at these lower levels provides reproducible insights to key proteins and metabolic pathway regulation (see below). These data and the reproducibility across nine repeat injections on five platforms (Figure 1 C) demonstrate how bottom-up proteomics data could be used alongside process monitoring, adding molecular information on chassis organism adaptation across a production run to add to current Critical Quality Attributes and Critical Process Parameters in industrial applications.

**Figure 1:**
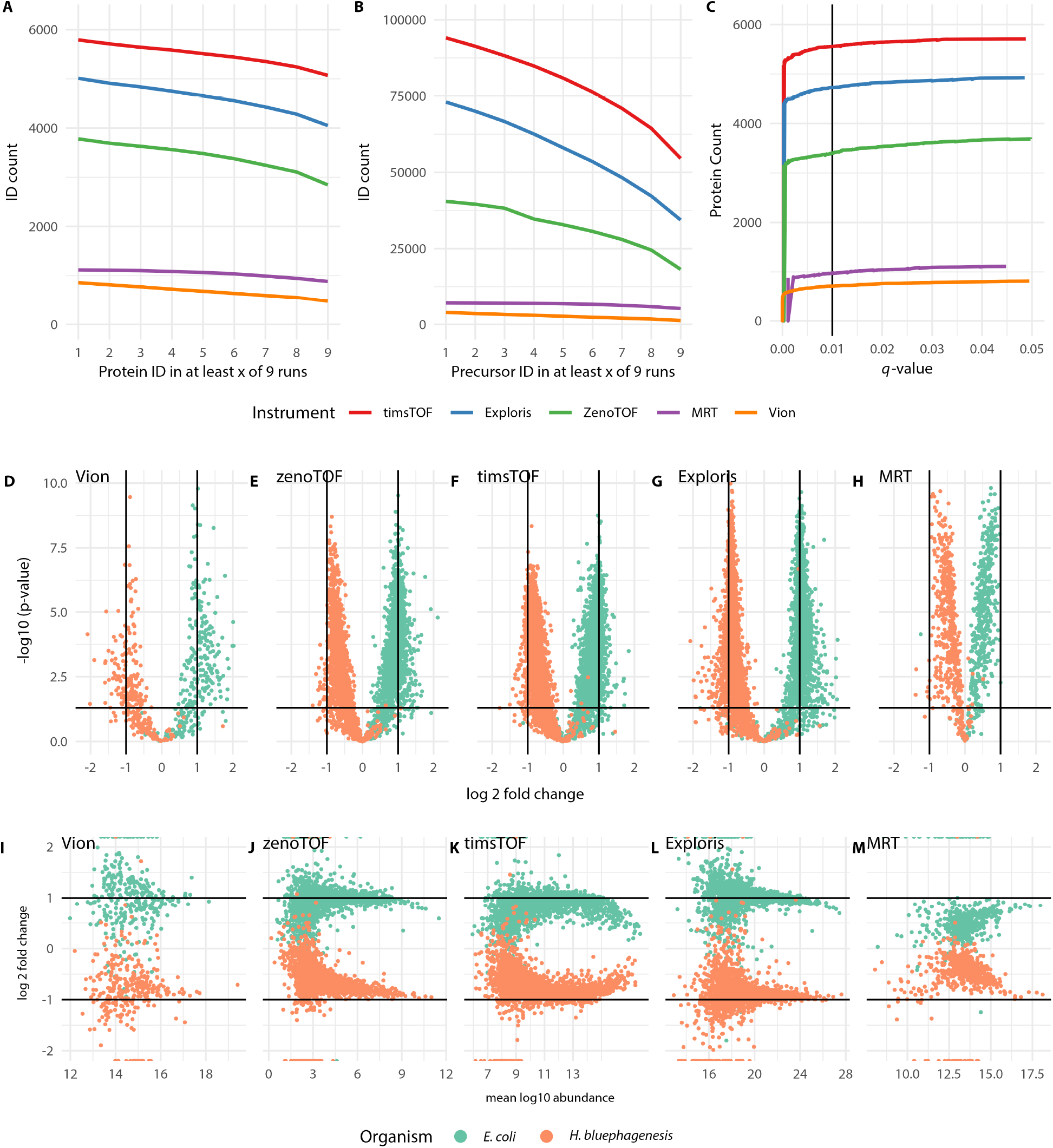
Consistency of protein (A) and peptide (B) identification across 9 injections. Protein identification *q*-value against number of proteins identified showing protein identification saturates around *q*-value 0.01, roughly a FDR of 1%. Volcano plots (D-H) and MA plots (I-M), both from left to right Vion, ZenoTOF, timsTOF, Exploris and MRT respectively, allow relative comparison of the instrument overall performance in terms of sensitivity to differential expression.

For comparison our previously published survey of *H. bluephagenesis* expressed proteins and assisted spectral library found 1160 proteins from a fractionated sample analyzed on a Waters XevoTOF^18^, where the most sensitive instruments here, the ZenoTOF, timsTOF and Exploris quantified 1758, 2928 and 2529 proteins respectively. In summary, substantially more of the proteome is covered on these instruments, than reported previously. Quantitative reproducibility at the peptide level Figure 1 B broadly aligns with the protein level data, with data from the more sensitive instruments (timsTOF HT and Exploris 480) providing a higher number of peptides per protein for quantification. Confidence in quantified protein identification is shown in Figure 1 C plotting number of proteins identified against *q*-value. The *q*-value is an estimate of the positive false discovery rate (pFDR) determined from the distribution of *p*-value when multiple statistical tests are made^20,21^. It is set so that the *q*-value is an estimate of the proportion of false discoveries for all tests with that, or a lower *q*-value. The rank order of instrument performance is consistent with the first two plots. Protein identifications in all cases come close to saturation before exceeding protein discovery *q*-value of 0.01.

Volcano plots (Figure 1 D-H and Figure S1 in supplementary information) and MA-plots (Figure 1 I-M), allow relative comparison of each instrument’s overall performance in terms of sensitivity to differential expression and quantitative accuracy. Volcano plots compare statistical confidence in calling a protein as differentially expressed against the estimated ratio of differential expression. In this case the “differential expression” has been set in the samples, so underestimate of protein expression, and higher *p*-values are likely caused by low signal to noise ratios for lower abundance peptides. MA-plots, report fold change estimates against mean abundance (Figure 1 I-M). All show a slight underestimation of ratio occurring for lower abundance, and so lower signal, proteins. Taken together these plots show how instruments sensitivity, signal-to-noise ratios and statistical performance in the detection of differential expression relate to each other. Although the total numbers of proteins quantified differs (as previously discussed), the quantitative accuracy of the four TOF-based instruments (timsTOF HT, ZenoTOF, MRT and Vion) and the Orbitrap-based Exploris 480 is similar (Figure 1 D-H). Pooling protein-level fold-change estimates from all instruments for the nominal 2:1 and 1:2 dilutions yielded a median observed fold change of 1.76 across all proteins identified in each organism with a median absolute deviation of 0.3, indicating modest ratio compression but overall good quantitative agreement on what we would predict (Figure 1 I-M). That this median measured ratio falls below the expected ratio of 2, is attributable to signal-to-noise ratios having a larger effect for low signal peaks. At higher signal the MA-plots also show a small underestimate of the expected ratio, it may be of course that small errors in quantifying peptide concentration or volume selection during sample preparation produced slightly lower than the intended ratio.

A couple of observations should be highlighted: The specified protein ratios were quantitatively detected with higher statistical confidence for data generated with the Exploris 480. Ratios from MRT data were compressed relative to expected values, which is likely due to background noise. We obtained early access to this platform as an example of a benchtop instrument. Further optimization is planned and results here represent the lower limits of that instruments’ capabilities. The superior depth achieved by the timsTOF HT and Exploris platforms reflects a combination of higher sequencing speed, improved ion utilization efficiency, greater dynamic range, and enhanced sensitivity for low-abundance peptides. The diaPASEF workflow additionally benefits from ion mobility-assisted precursor separation, improving spectral complexity management and duty cycle efficiency^22^. In contrast, lower price point or legacy platforms exhibited reduced proteome depth, likely reflecting limitations in sensitivity, ion transmission efficiency, and precursor separation capacity rather than deficiencies in quantitative reproducibility itself. PAXdb is a curated protein abundance database that integrates experimentally derived estimates of absolute protein abundance across organisms and conditions. PaxDb provides a community resource which maps re-processed publicly available protein expression data to a single common namespace (STRING^23^) thus providing a summary of current knowledge of protein expression in organisms such as *E. coli*, and specific tissues and organs of more complex organisms. We used it here as an orthogonal benchmark for assessing whether relative abundance trends observed in our datasets were biologically plausible. Direct comparison of the absolute abundance of the *E. coli* proteins from each instrument against each other, and the PaxDb database of absolute protein abundance (https://pax-db.org/species/511145)^24^ is reported in the supplementary information (Figure S2). Comparison of protein abundance against the PaxDb provides a reference point for these instruments compared to expected protein expression. The highest correlation was observed between the Exploris 480 and the ZenoTOF (0.898), with reduced consistency apparent between the Exploris 480 and timsTOF HT (0.873). More importantly perhaps, particularly in the context of systems modelling, absolute abundance correlated relatively well (>0.65) between the timsTOF HT, Exploris 480 and ZenoTOF with data from PaxDb. It is clear from this comparison that quantification of low abundance proteins is somewhat compromised with both the Vion and MRT instruments, explaining the reduction on the overall numbers of quantified proteins. Additionally, the three higher coverage instruments (timsTOF, Exploris and ZenoTOF) overestimate the higher abundance proteins with respect to the PAX database values.

### Biological Insight

An important output from monitoring protein abundances in biomanufacturing chassis organisms is how these can be used to better understand changes in key metabolic pathways. The percent protein coverage of key KEGG pathways in *E. coli* achieved with each instrument is reported in Figure 2 for pathways with 50 or more associated proteins. Proteins from each pathway that were identified by all instruments are found in supplementary dataset 1. All instruments can report across a large number of pathways with critical biological function. As expected from the depth of proteome coverage, the timsTOF HT, Exploris 480 and ZenoTOF consistently deliver the highest coverage. The most important pathway for understanding carbon flux, critical to biomanufacturing, is carbon metabolism, for which even the Vion achieved 43% coverage, compared with 91% for the timsTOF HT. Obtaining quantitative data for almost half of this pathway is still useful and an example of how a lower cost/sensitivity platform such as the Vion, can provide sufficient information of importance to a biomanufacturing facility. A corollary is the relatively low (<10%) coverage of the ABC transporters, for which over 80% is covered by the timsTOF HT. Such transporters could be critical in a given biomanufacturing chassis for their role in the export of toxic product molecules to media; understanding and optimizing their contribution will necessitate deeper proteome profiling^25,26^.

**Figure 2:**
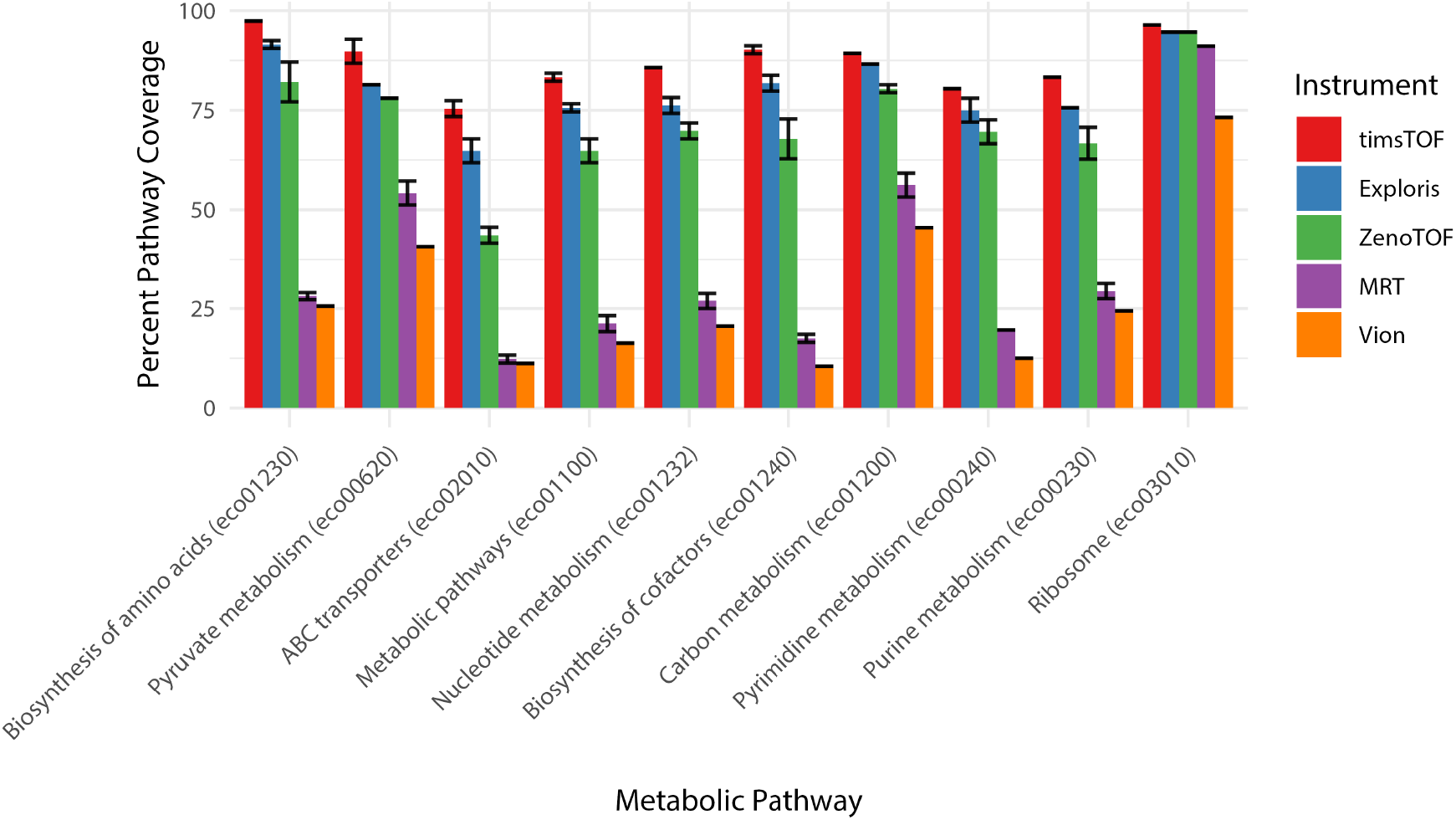
For the data obtained from *E. coli* the median percentage coverage of proteins annotated in KEGG as belonging to each of the annotated pathways per data file, with median absolute deviation in identification numbers plotted as error bars.

### Exemplar Applications

#### Alternate Carbon Feedstocks

As an example of the power of proteomics to provide insight into the biology of novel chassis organisms for biomanufacturing, we used the ZenoTOF to explore changes in the proteome of *H. bluephagenesis* cultured in two different sources of carbon: glucose which enters its core metabolism via the glycolytic pathway, or acetate which enters via conjugation with Coenzyme A (Figure 3). The ZenoTOF was selected for this carbon-source exemplar not because it gave the deepest proteome coverage, but because it represents a mid-tier platform that provided sufficient coverage for biologically interpretable pathway analysis. Using this approach we were able to quantify 2,818 proteins from the 3,821 entries in the *H. bluephagenesis* uniparc proteome (UP000838464). Of these, 120 proteins (4.3% of those quantified) were statistically significantly differentially regulated, with 58 being elevated and 42 being at lower levels (considering those at an adjusted *p*-value <=0.01, of fold-change >2). The proteins names from the uniparc *H. bluephagenesis* sequence databases were mapped to KEGG gene names and from this to KEGG KO numbers which indicate gene function. A volcano plot (Figure 3) shows the significance of differentially expressed proteins with KO numbers linked to KEGG pathways hbp00010 (Glycolysis / Gluconeogenesis) and hbp00620 (Pyruvate metabolism), the latter of which includes metabolism of acetate including its conversion to acetyl-CoA and so the direction of carbon flux into the citrate cycle. A protein annotated with K01895 (acetyl-CoA synthetase) is upregulated in acetate fed culture over glucose fed culture. This is to be expected as the bacteria must compensate for loss of glucose by increasing carbon flux from acetate directly into the citrate cycle. Another set of proteins with KEGG annotations K01571, K00845, K00134, K00873, K01810 are all down regulated in acetate fed cultures with respect to glucose fed cultures, are part of the glycolysis pathway. It follows that growing *H. bluephagenesis* in an environment with reduced glucose will diminish expression of proteins required to make use of it. That these proteins belong to pathways that we might expect to be modulated under the respective feedstock conditions, provides confidence in the general utility of our quantitative proteomics data for such interrogations. Further research into how and why the levels of these proteins are altered when *H. bluephagenesis* is grown in different carbon sources will provide insights to how any novel chassis system under active development for biomanufacturing applications could be controlled and adapted. Understanding how such organisms respond to changing conditions will guide selection of culture conditions. In the specific case of feedstocks, which are a major cost in biomanufacturing, understanding and modifying organisms capability to grow on various feedstocks may help transitions a biomanufacturing platform to a lower cost carbon source^27,28^. Understanding the promoters driving differential expression of these proteins under differing feed stock conditions guide selection of promoters for use in production pathways, and even enable initiation of production by switching feed stock. We are currently exploring genetic interventions to boost, or suppress, expression of proteins of interest identified in this way could increase yield of products in model bioreactors. This demonstrates how relatively straightforward quantitative proteomics experiments can produce useful information to guide development for biomanufacturing production.

**Figure 3:**
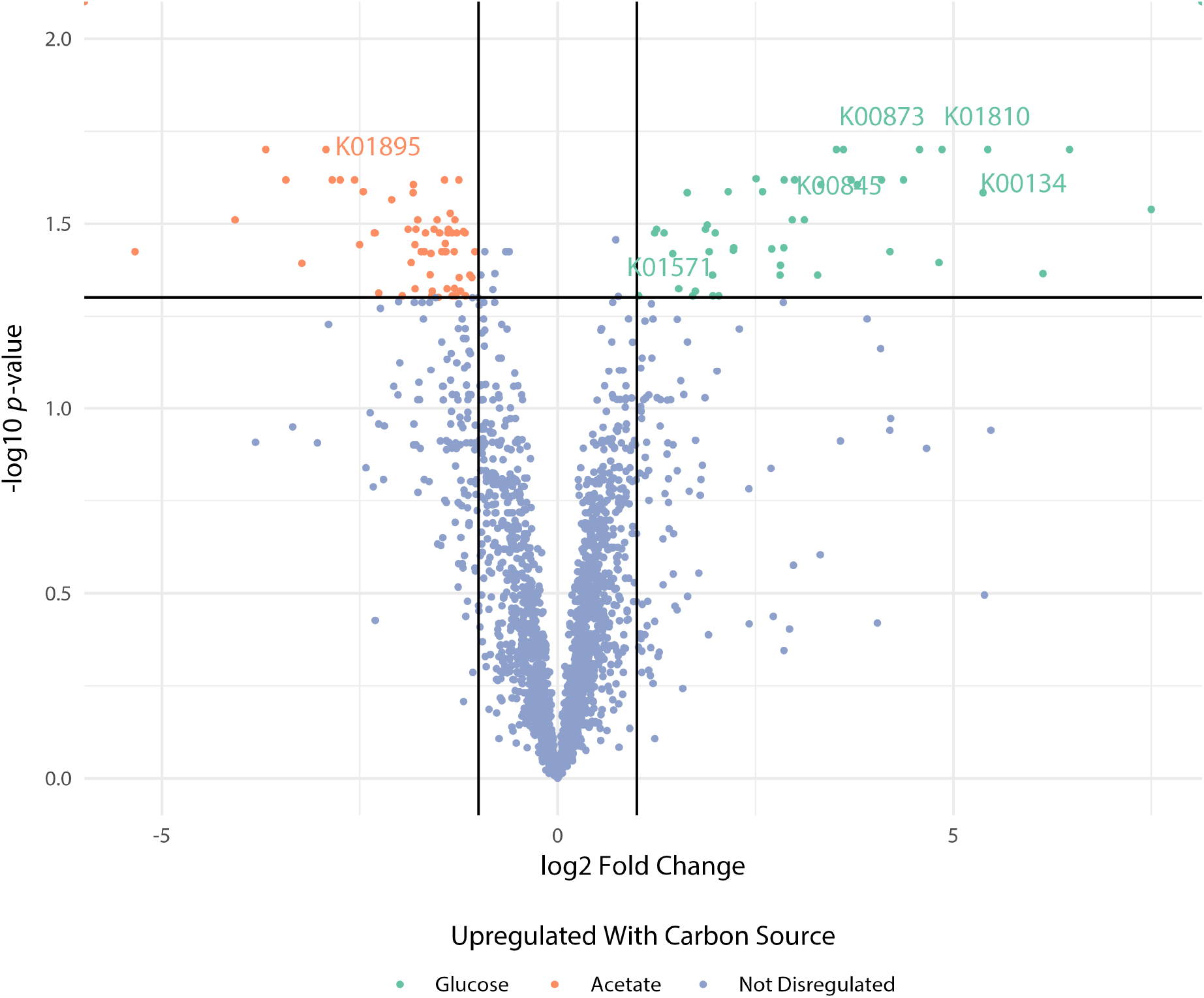
Volcano plot showing protein dysregulation in *H.bluephagenesis* grown with carbon sources either glucose or acetate from proteomics data acquired on Sciex ZenoTOF. Proteins with statistically significant (adjusted *p*-value <=0.01) dysregulation greater than 2-fold between conditions are highlighted, and dysregulated proteins associated with glycolysis or acetate metabolism are annotated with Uniprot accension numbers.

#### From Discovery to Targeted Proteomics: Confirming Vitamin B12 Biosynthesis in *H. bluephagenesis*

Having demonstrated the utility of global DIA proteomics for characterizing novel chassis organisms, we next explored a transition from discovery to targeted proteomics — a workflow that we have previously suggested as practical biomanufacturing applications. In such experiments, peptides from candidate proteins identified through high-end DIA experiments and found to report on critical metabolic processes are subsequently monitored using lower-cost targeted instruments. Discovery datasets such as those presented here, would both map the pathways and identify which peptides are suitable for targeted MS quantitative analysis. Such experiments could be used for routine quality control and process monitoring at production facilities.

As a case study, we investigated the vitamin B_12_ biosynthesis capability of *H. bluephagenesis* (Figure 4). Establishing endogenous B12 production was important because ongoing engineering efforts aim to introduce B_12_-dependent production pathways into this chassis. *H. bluephagenesis* was able to grow in media free of exogenous B_12_ — suggesting endogenous synthesis. For any relatively underexplored organism it is useful to confirm putative functional gene annotations and here this pathway was chosen as an exemplar.

**Figure 4:**
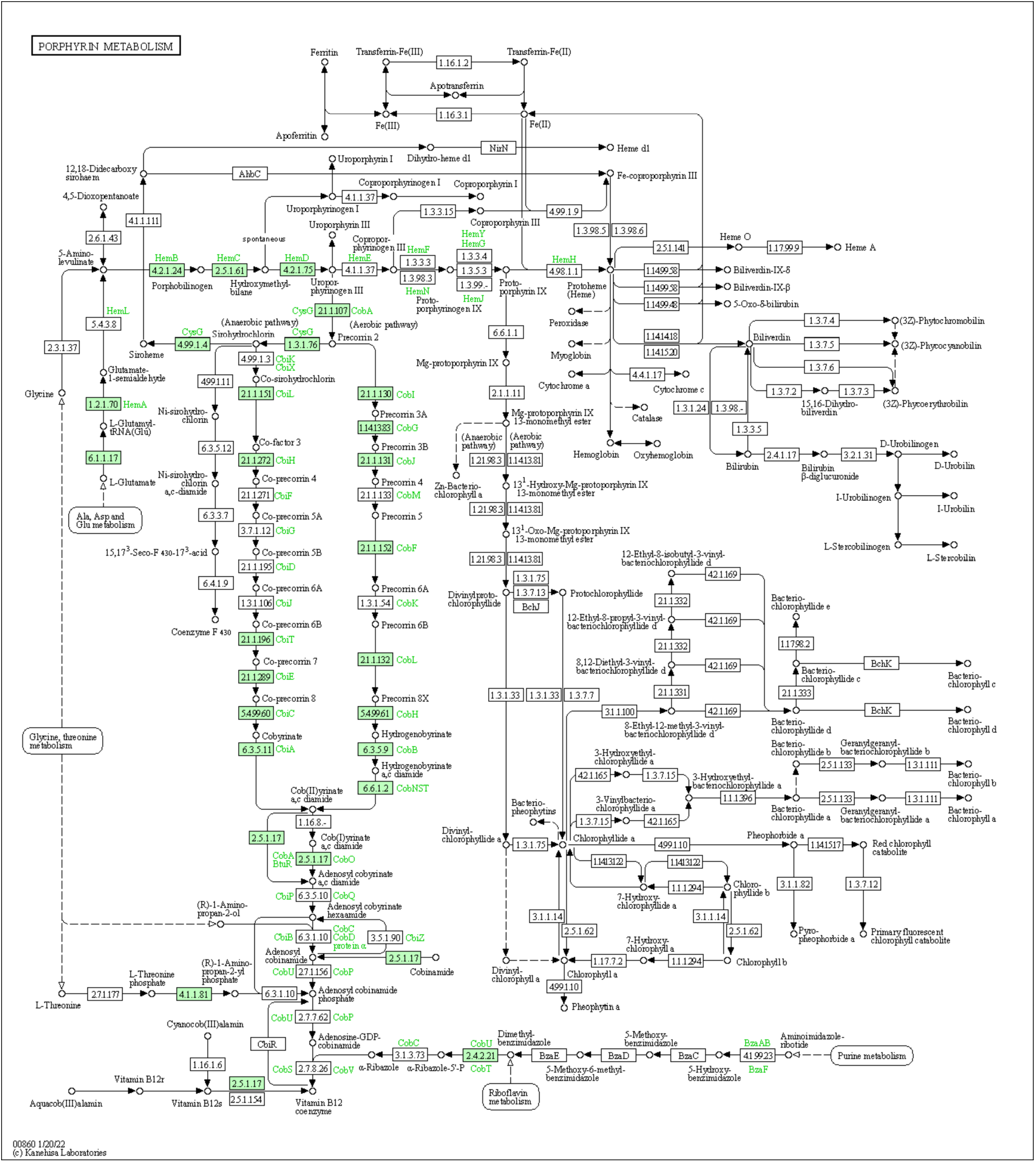
Protein sequences with peptides identified by targeted proteomics analysis of H. bluephagenesis’s putative endogenous B_12_ producing pathway projected onto to Kegg “Porphyrin metabolism” pathway. Proteins confirmed as present in the sample are colored green.

Using the genomic sequence of *H. bluephagenesis*, we identified the proteins annotated as components of the vitamin B_12_ biosynthesis pathway (Figure 4). Diagnostic peptides unique to each candidate protein were selected and synthesized as unlabeled standards. These reference peptides were used to develop targeted assays, which were then applied to whole-cell lysates of H. bluephagenesis to confirm the presence of the corresponding proteins. The detected proteins were mapped onto the KEGG vitamin B_12_ biosynthesis pathway (Figure 4), achieving coverage of the majority of pathway steps (Figure 4). This provides strong evidence that H. bluephagenesis possesses a functional B12 biosynthesis pathway, supporting its suitability as a host for B12-dependent engineered pathways.

This approach illustrates a practical workflow we envision for biomanufacturing proteomics. Initial discovery-phase experiments, performed on high-specification instruments can identify proteins of interest — whether components of an engineered production pathway, indicators of organism viability, or markers of metabolic state. Peptides from abundant and accessible candidate proteins from these pathways are then translated into targeted assays deployable on substantially lower-cost platforms, such as triple quadrupole mass spectrometers, which are well suited for yes/no quantitative answers using selected reaction monitoring (SRM) or multiple reaction monitoring (MRM)Such instruments could be employed at production sites, providing routine confirmation that engineered pathways are actively expressed, and monitoring key metabolic indicators as part of a quality control framework.

## Discussion

Microorganisms based biomanufacturing faces key challenges^29^. Better control of established organisms is required to improve yields and cut costs, while accelerating adoption of lesser-known species with valuable traits. Our investigations model these scenarios using proteins from established *E. coli* alongside recently isolated *H. bluephagenesis* sp TD01^5,7,8,29–33^.

*E. coli* is the dominant biomanufacturing organism by historical accident^34,35^. Despite a knowledge base from over century of research on this model organism, implementing novel engineered biology pathways in *E. coli* remains difficult^28^. While some alternative species see limited use (Bacillus subtilis^36^, Streptomyces^37^, Pseudomonas putida^38^, Corynebacterium glutamicum^39^), nature likely contains species with more favorable traits. These include toxicity resistance, efficient carbon utilization, and an ability to thrive in economical bulk production conditions^40,41^. *H. bluephagenesis* demonstrates such advantages, it thrives in high-salt, high-pH environments hostile to most bacteria, enabling cost reducing non-sterile cultivation^42^. Here we have demonstrated how semi quantitative discovery proteomics can assess the metabolic processes that occur in proposed biomanufacturing strains and how such insights can determine how a given organism adapts to changes in its energy source or to follow the effectiveness of a given pathway. These data could guide subsequent engineering strategies by identifying bottlenecks in carbon utilization, redirecting metabolic flux toward desired products, or selecting process conditions that favor target pathway activity.

For biomanufacturing to replace petroleum-based industry, it must produce commodity chemicals at competitive costs, which can only be achieved when there is understanding of the pathways that can be optimized for maximal yield. Effective proteomics support for biomanufacturing requires methods and platforms spanning development through production. Building on prior DIA benchmarking studies^17^, we have tested five different MS platforms, which are representative of what might be available to small medium and large entities for their utility. By comparing data from tryptic digests of proteins from *E. coli* K-12 MG1655 and *H. bluephagenesis* in 1:2, 1:1, and 2:1 ratios, we assess coverage, sensitivity, quantitative reproducibility and accuracy. We demonstrate here that affordable MS instruments that yield partial proteome coverage can still provide sufficient data to support investigation of proteome-level changes with a view to optimization of economically viable production. Targeted monitoring of these proteins could subsequently provide an operational readout of metabolic state during scale-up or process optimization.

While higher-sensitivity instruments were able to quantify greater proportions of the two bacterial proteomes investigated here, lower-sensitivity workhorse platforms could still be used effectively to monitor core metabolic pathways. While high-end platforms such as timsTOF and Exploris provide the greatest depth, useful process-relevant biological information can also be obtained on more accessible instruments, and this is why we demonstrated the effects of altering carbon source with the ZenoTOF, representative of mid-price range instrumentation. Such instruments could therefore find a place in troubleshooting or process control, such as the FDA/EMA Quality by Design (QbD) regulatory framework for biopharmaceuticals^43–45^, relating key metabolic pathways to bioreactor productivity and product quality. Alternatively, instruments that enable more in-depth and robust protein quantification may be better suited for initial discovery studies as a pre-cursor to targeted, analyte focused assays using selected/multiple/parallel reaction monitoring (SRM/MRM/PRM). A use case for targeted proteomics outlined above in which the presence of the co-factor B_12_ production pipeline in *H. bluephagenesis* was confirmed by MRM. This confirms that the availability of B_12_ in the *H. bluephagenesis* chassis required for a particular production pathway. Such an approach might easily be adapted for QC approaches to monitor bioreactors to confirm continuing the health and productive capacity of a production culture, drawing on prior development work using global DIA approaches to inform selection of a small set of proteins for targeted analysis. We have demonstrated the utility of such approaches previously for over 1,800 proteins in *S. cerevisise*^46^ and for biomarkers of health in CHO cell culture^14^. Where targeted proteomics by mass spectrometry is not available non-MS methods such as qPCR or multiplexed immunoassays such as developed in health care settings^47^ could be used for QC informed by discovery phase proteomics.

The instrument systems studied here demonstrate a wide range of performance capability. In terms of depth of proteome coverage both the Exploris and timsTOF performed similarly capturing close to the complete proteome. These instruments have very high scan rates but operate on differing principles. The timsTOF accompanies this scan rate with gas phase separation of peptides by ion mobility, enabling sequential fragmentation and detection of peptides ordered by their length, this minimizes interference between analytes, and other background noise. The Exploris isolates narrow windows of peptides by mass, in a manner similar to SWATH, described below, and sequentially passes these via fragmentation to an orbitrap mass analyzer, which has a high resolution able to resolve signals from each other and also from noise. The zenoTOF performed less well in terms of depth of coverage than the instruments described above, although still quantified in excess of 3000 proteins across eight of nine injections, which compares favorably with a single bacterial genome coding for about 4000 proteins. The zenoTOF performs DIA in a similar manner to the Exploris, by isolating peptides within narrow bands of m/z in a trap, then passing them as a pulse to the ToF mass analyser. It may be that this SWATH dictated slower scan rate and the slightly lower resolution of ToF over orbitrap accounts for the lower depth of coverage of the two top performing instruments. The Waters MRT is an altogether different type of instrument to the three described above. It has no precursor separation, rather flips between precursors and their fragments, however it’s multipass TOF enables very high-resolution mass spectra, which should separate ions of very similar mass thus seeing through noise and mutual interference. The “desktop” feature of the MRT could be attractive in biomanufacturing environments. While high-end platforms such as timsTOF and Exploris provide the greatest depth, useful process-relevant biological information can also be obtained on more accessible instruments, and this is why we demonstrated the effects of altering carbon source with the ZenoTOF, representative of mid-price range instrumentation.

Our study has limitations. LC configurations were not identical across all platforms, acquisition methods were not exhaustively optimized for maximal depth, and one platform required an alternative processing workflow. Accordingly, the results should be interpreted as representative comparisons of deployable workflows rather than definitive benchmarks of absolute instrument capability. Despite this, we demonstrate that quantitative proteomics can provide crucial insights during production process development, particularly with novel organisms, and could further support production monitoring and troubleshooting. Whilst this work was not intended as an in-depth mechanistic study of *H. bluephagenesis* metabolism, the biological experiments included as exemplar applications demonstrate how DIA proteomics can support chassis characterization and process-informed monitoring. Within that scope, the acetate/glucose comparison identifies expected and interpretable changes in carbon metabolism, including acetyl-CoA synthetase and glycolytic pathway components, while the vitamin B_12_ work uses targeted peptide assays to confirm expression of proteins across a pathway of practical relevance to ongoing engineering work. Collectively, these findings support a tiered model for proteomics deployment in biomanufacturing. High-end DIA platforms provide deep systems-level insight during strain development and process optimization, while lower-cost or even ‘legacy’ platforms remain capable of monitoring biologically informative subsets of the proteome relevant to process control and troubleshooting. We therefore envision proteomics becoming an increasingly integrated component of engineering biology workflows, spanning early-stage discovery through to operational deployment.

## Methods

### Bacteria Strains and Cultures

#### Halomonas bluephagenesis sp

TD01 and *E. coli* (K12/MG1655) samples were processed identically. LB Broth Miller was prepared from powder (Formedium, 25 g L^−1^ comprising tryptone 10 g L^−1^, yeast extract 5 g L^−1^ and sodium chloride 10 g L^−1^, pH 7.0) and for *H. bluephagenesis* adjusted to 5% sodium chloride (additional 40 g L^−1^) and pH adjusted with sodium hydroxide (5 M, 4 mL L^−1^, to pH 9). Liquid media was sterilized by autoclave. For plates, agar (15 g L^−1^) was added to the appropriate sterile lowor high-salt media described above and heated in a microwave (30% power, 20 min) then poured into plates and allowed to cool in a laminar flow hood. Plates were stored (5 °C) for up to four weeks prior to use.

Cells from a glycerol stock were aseptically inoculated onto LB-agar plates, incubated (37 °C, overnight) then transferred to cold storage (4 °C). Single bacterial colonies of *E. coli* or *H. bluephagenesis* were inoculated aseptically into liquid media (50 mL) in a 250 mL conical flask. Cultures were incubated (37 °C, 200 rpm) and OD600 monitored. When OD600 reached 0.3–0.6, in which growth was late log/early stationary phase, culture aliquots (450 µL) were recovered to pre-weighed 5 mL microfuge tubes on ice and pelleted by centrifugation (3200 g, 10 min, 4 °C). Supernatant was discarded and the wet mass of the pellet determined by difference with tubes empty mass. Pellets were frozen (−20 °C) for later lysis and digestion.

To determine the effect of carbon feed stocks glucose or acetate were prepared from *H. bluephagenesis TD1.0* (a T7-like RNA polymerase-integrated derivative of *H. bluephagenesis sp TD01*^48^) cultures grown in 100 mL MM63 supplemented with 5 gL^−1^ glucose or acetate and harvested in the exponential growth phase after OD600 exceeded 1. Cultures were pelleted by centrifugation (3200 g, 10 min, 4 °C), followed by three washes with phosphate buffered saline and repeat centrifugation. Pellets were frozen (−80 °C) for later lysis and digestion.

#### Pellet Recovery and Digestion

Pellets were simultaneously defrosted and resuspended in S-trap^49^ lysis buffer (7% SDS, 70 mM TEAB, pH 7.55, 3 µL buffer per mg wet mass of pellet). DNA was fragmented by addition of benzonase (Sigma/Merk, 1 µL) and sample vortexed briefly. If required, a second addition of benzonase removed any remaining viscosity. Cell debris were removed by centrifugation (4200 g, 2 min). Samples were buffer exchanged by ultrafiltration (Amicon Ultra 0.5 mL 3 kDa cut-off, 4 washes) with further SDS-free lysis buffer (70 mM TEAB, pH 7.55) removing lipids, carbohydrates, nucleic acid fragments and salt.

Protein was digested with the S-trap^49^ mini kit (Protifi) according to the manufacturer’s instructions, the sample already being in the recommended SDS-lysis buffer (see above). Briefly, sample volume was taken to give 1–5 mg total protein, reduced with TCEP (60 °C, 30 min, 50 mM final concentration), alkylated with iodoacetamide (room temperature in the dark, 30 min, 40 mM final concentration). The sample was acidified with phosphoric acid (1/10 vol, 1.2% final concentration) and diluted in S-trap binding buffer (7 vols, 90% methanol, 100 mM TEAB). The sample was loaded onto S-trap in aliquots of 190 µL aliquots followed by centrifugation (4,000 g, 1 min). The S-trap was washed with S-trap binding buffer (4 × 150 µL) with centrifugation as for loading. Protein was digested by addition of trypsin (Promega, 20 µL, 0.5 µg µL^−1^) in digestion buffer (50 mM TEAB) and incubated (47 °C, 1 hr) in a water bath.

#### Peptide Recovery Estimate

The yield of peptides was determined by the Pierce quantitative fluorometric peptide assay (PN:23290) according to the manufacturer’s instructions. Fluorometry was performed on a CLARIOstar Plus plate reader (BMG Labtech) and data processed with Mars v3.40 R2 fitting a four-parameter curve.

#### High pH Fractionation for Vion Spectral Library

To enable analysis of HDMSe data obtained on the Vion, a spectral library was acquired from a set of fractionated samples. Peptides (50 µg, 300 µL 0.1% TFA) were loaded onto a high-pH C18 spin column (Pierce peptide desalting columns PN:89852) each column and centrifuged (3000 *g*, 2 min), washed with pure water (300 µL) and centrifuged as above. Peptide was eluted in fractions by stepwise addition of stepwise increasing acetonitrile concentration (5, 7.5, 10, 12.5, 15, 17.5, 20, 50% acetonitrile, 300 mM ammonium bicarbonate adjusted to pH 10 with ammonium hydroxide), collecting as separate fractions. Fractions were lyophilized by vacuum centrifugation. Peptide was assumed to be approximately equally split across fractions giving 6-7 µg per fraction.

#### Fixed Ratio Samples

The fluorometric estimate of peptide recovery from both the *E. coli* and *H. bluephagenesis* whole cell lysates allowed the concentrations of both to be equalized at 0.2 µg µL. Volumes of *E. coli* and *H. bluephagenesis* whole cell lysates (150:150, 200:100, and 100:200 µL) were combined to give samples of 300 µL, 60 µg total peptide in species specific ratios of 1:1, 1:2 and 2:1 in a similar approach to that adopted in multi-site comparison studies. Each sample was divided into aliquots (10 × 30 µL, 6 µg each) and lyophilized alongside aliquots of pure single species peptide (30 µL, 6 µg each).

Further details on the chromatography and acquisition conditions for each of the instruments deployed is found in the Supplementary information.

### Targeted Proteomics

#### Standard Peptide Selection

Protein sequences were identified by homology in the TD1.0 genome. Candidate peptides were selected by importing the proteome file into Skyline; *in silico* digest and peptides filtered to those between 8 and 20 amino acids in length; excluding the first 25 N-terminal amino acids; excluding potential ragged ends;and excluding peptides containing methionine; arginine-proline or lysine-proline motifs. Two of the small target proteins would not produce tryptic peptides suitable for analysis by LC-MS, these were: Adenosylcobyric acid synthase; and Cob(II)yrinic acid a_c-diamide reductase. The remaining protein sequences suggested several of possible target peptides per protein. These peptides were purchased from JPT Peptide Technologies GmbH peptides as unpurified, unlabeled sequences.

#### Pellet Lysis and Tryptic Digestion

Following the same pellet recovery procedure as above the protein was digested with the S-trap mini kit (Protifi). Prior to digestion the total protein quantification was determined by the Pierce BCA assay (Thermo Scientific). Sample volume was taken to give 150 µg total protein which was reduced with TCEP (50 mM final concentration, 60 °C, 30 min), alkylated with iodoacetamide (40 mM final concentration, room temperature in the dark, 30 min). The sample was acidified with phosphoric acid (1/10 vol, 12%) and diluted in S-trap binding buffer (7 vols, 90% methanol, 100 mM TEAB). The reduced and alkylated sample was then split between two S-traps for a total of 75 µg protein load per trap; in cycles of 190 µL aliquots followed by centrifugation (4,000 × g, 1 min). The S-trap was washed with S-trap binding buffer (4 × 150 µL) with centrifugation as for loading. Protein was digested by addition of trypsin (Promega, 20 µL, 0.5 µg µL^−1^) in digestion buffer (50 mM TEAB) and incubated (47 °C, 1 hr) on a heated block. Peptide was recovered by washes of digest buffer (50 mM TEAB, 40 µL), aqueous formic acid (0.2% formic acid, 40 µL) and organic buffer (50% acetonitrile, 0.2% formic acid, 40 µL) with centrifugation as for loading above. The peptides were lyophilized by vacuum centrifugation, resuspension (30% acetonitrile, 1% formic acid, 50 µL), and further vacuum centrifugation. Post digestion the peptide concentration was verified by Pierce Peptide Fluorescent Assay (Thermo Scientific) to verify load on column.

#### LC-MS Analysis

MRM analysis was conducted on a Waters Acquity H-class UPLC system coupled to a Waters Xevo TQ-S triple quad mass spectrometer. The LC system was set up with a mobile phase A of HPLC grade water and mobile phase B of HPLC grade acetonitrile (both Honeywell LC-MS Chromasov) both acidified with 0.1% formic acid. Sample was injected onto column (Acquity UPLC Peptide CSH C18, 1 mm ID, 150 mm length, 1.7 µm particle size, 130 Å Pore Size), held at 50 C, with 3% B and a flow rate of 0.1 ml/min. Peptides were eluted with a linear analytical gradient of 3-40% over 30 min. The column was then washed by ramping up to 90% B over 1 min then holding for 4 min. Finally the system was re-equilibrated at starting conditions of 3% B for a further 5 min.

MRM method setup was and data analysis was mediated through Skyline v22.2.0.351. A stepwise approach was taken first to determine retention time of each of the standard peptides; then to optimize the collision energy for each transition using Skyline’s built in optimization feature; and finally to export scheduled MRM methods. During this process peptides that gave poor signal were excluded from the analysis. The dwell time per transition was set at 30 ms, the time window around expected retention time was set to 2 min, the maximum concurrent transitions was set to 50. Two methods were required to cover the full set of peptides. Peaks were typically 30 s at base with a minimum of 19 scans across the elution profile.

For MRM method development standard peptides were injected at approximately 50 pmol/peptide/injection. For final analysis sample was injected with 10 µg total peptide per injection.

The batch set up for the final analysis comprised a blank wash injection; blank injections to confirm minimal carry over from standards or previous injection; then sample injection. Four replicate injections were carried out to confirm the result.

#### Data Processing DIA

The complete data processing pipelines were scripted for command line tools and both the script and an explanatory pdf including both code and documentation are included in supplementary information. Brief descriptions of each processing pipeline are included in the supplementary information.

#### Statistical Analysis DIA

All data analysis was performed in R version 4.6.1 (2026-06-24 ucrt), data sets were quantified and differential expression between samples calculated using MSStats^50^. Scripts enabling reproduction of the analysis are included in supplementary information and with the data deposition at PRIDE^51^ supporting this publication.

#### Data Processing Targeted Analysis

The final data set was imported into Skyline. Each peptide and its transitions were visually interrogated to check the peak in the sample closely matched the standard peak and was absent in the blank injection. Peptides that were not confidently identified were removed, and proteins without remaining peptides were also removed. Proteins with confidently matched peptides were mapped to KEGG pathways blastkoala.

## Supporting information

Supplementary data set 1

Supplementary data set 2

Supplimental Information - Documented Scripts

## Supplementary Information

The supplementary information contains further experimental details pertaining to acquisition parameters for each instrument used. Processing pipelines are detailed. In addition, it contains a zoom of the volcano plots in Figure 1 main text, with wider x-axis and in larger format to facilitate comparison and also plots of log_10_ protein abundance estimates from MSstats analysis of *E. coli* data from each of the instruments and the PaxDb data base.

## Data Availability

Further data referred to in this manuscript are found in the supporting data including excel files with a list of all peptides targeted and proteins found as shown in the figures. All Raw mass spectrometry data is available via ProteomeXchange with identifier PXD071869^52^.

## Acknowledgements

This work was supported by the Future Biomanufacturing Research Hub (grant EP/S01778X/1), funded by the Engineering and Physical Sciences Research Council (EPSRC) and Biotechnology and Biological Sciences Research Council (BBSRC) as part of UK Research and Innovation. Instrumentation was funded by BBSRC (grants BB/W019892/1 and BB/T018127/1). We acknowledge the support of staff in the Mass Spectrometry and Separations Facility of the Faculty of Science and Engineering at the University of Manchester and in the Centre for Proteome Research (CPR) at the University of Liverpool. Finally, we thank Waters for their continued support to the Michael Barber Centre for Collaborative Mass Spectrometry and specifically Lee Gethings and Dale Shepherd for their help in acquiring data on the MRT.

