## Supplementary material for "From Discovery Proteomics to Process-Informed Monitoring in Biomanufacturing Chassis Development": Supplimental Information


#### Supplementary Methods

##### LC-MS Waters Vion

The full sample set now comprised eight fractions each of *E. coli* and *H. bluephagenesis* peptides to generate a spectral library on the Vion; combined ratios of whole cell lysate digests at 1:1, 1:2 and 2:1 to assess instrument performance and a pure species specific sample for each species, each sample of which aliquoted at of 6 µg lyophilized peptide. For LC-MS analysis lyophilized peptides were resuspended in loading buffer (3% v/v acetonitrile, 0.1% v/v formic acid, 24 µL, 0.25 µg µL$^{-1}$ final peptide concentration). The fractionated samples were injected onto the Vion, in order to create an instrument specific spectral library.

Data was acquired in triplicate for 1:1, 2:1, 1:2 and single species samples on each of the instruments in the study.

#### Liquid Chromatography

Peptide were injected onto a Waters Acquity M-Class with a nanoEase M/Z peptide CSH 130 Å pore size, 1.7 µm particle size 150 µm ID, 150 mm in length in a column oven at 50 °C at starting conditions of 97% mobile phases A (10.1% formic acid) and 3% mobile phase B (acetonitrile, 0.1% formic acid) at 2 µL $min^{-1}$ for three minutes. Peptides were eluted with an analytical gradient of 3-30% B over 3-65 minutes, followed by a steeper gradient to 45% B by 78 minutes. The column was then washed with 95% B for 5 minutes and equilibrated to starting conditions.

#### Vion HDMSe

LC-eluent entered the Vion IMS QTOF (Waters) instrument via the Waters nanospray source in positive polarity and sensitivity analyser mode. Nebulising gas flow to the emitter was supplied directly from the gas line supplying the instrument through a control valve, rather that by the instrument itself. Data was acquired in HDMSe mode m/z range 120-2000 with 1 s scan times. Low energy scan collision energy was 7 V, high energy collision energy ramped from 22-28 V. Voltages and gas flows were as follows Capillary: 4 µV, Sample cone voltage 40 V, Source offset voltage 80 V, Cone gas: 50 L $h^{-1}$; Purge gas: 350 L $h^{-1}$.

#### LC-MS Sciex ZenoTOF

##### Liquid Chromatography

Peptide were injected onto a Waters Acquity M-Class with identical setup and method to that described for the Vion above.

#### ZenoTOF SWATH-MS

LC-eluent entered the ZenoTOF 7600 (Sciex) mass spectrometer through a Sciex 1-50uL Micro/Microcal optiflow source with a 1-10 µL emitter installed. The source temperature was 225°C and gasses were as follows, curtain gas 35; CAD gas 7; ion source gas 1 20; and ion source gas 2 60 PSI The SWATH-MS method comprised an MS-TOF scan (0.1 s accumulation time; m/z range 400–2050 ) followed by 100 Zeno-trap SWATH scans (0.018 s accumulation time; m/z range 100-2000) of variable widths with stepped collision energy (see supplementary data for tabulated values). Data was acquired for minutes 8-35 post injection in a 50 minute method, the total scan time was 2.394 s and 1253 complete scans were acquired through the acquisition.

#### Exploris 480

##### Liquid Chromatography

Peptide were loaded onto Evotips using the manufacturers protocol and analysed by the Evosep One LC system (Evosep) using the inbuilt "15 samples per day" method. The operation of the EvoSep is different to that of the other LC systems in the study, but for comparison the analytical gradient approximates to 100% mobile phase A at 220 nL min$^{-1}$ to 35% B over 88 min through a Thermo Scientific ES906 Easyspray column with built in emitter (C18 150  m x 150 mm 2  m column).

##### Exploris DIA

Data was acquired on an Exploris 480 mass spectrometer (Thermo Scientific) in DIA profile mode. The method comprised 42 DIA fragment window scans with a full scan every 14 scans, for a total of 45 scans per duty cycle. Full scans were acquired in the m/z range 345-900 at 120,000 resolution with 300% normalised AGC target and automatically selected max IT.

Data-independent fragments were acquired in 42 13 m/z fragmentation windows selected with “window placement optimisation” without overlap; over an acquisition range 350-896 m/z; with 15000 resolution; NCE at 32%; and AGC target set to 1000% A full scan was acquired after every 14 DIA scans, so it took 3 full scans (with each CV) to fragment the ions in all 42 DIA windows.

#### timsTOF HT

##### Liquid Chromatography

Peptides were injected onto a NanoElute (Bruker Daltonics) ultra-high-pressure Nano-flow chromatography with a trapping column (Thermo Trap Cartridge, PepMap100, C18, 300 m x 5 mm) and analytical column (PepSep, twenty-five Series 150 µm, 1.5 µm column). Peptides were eluted with an analytical gradient of 2-35% B over 90 minutes at a flow rate of 0.5 L/min, then washed at 95% B before re-equilibration to starting conditions.

##### TimsTOF DIA

Data was acquired on a TimsTOF HT mass spectrometer (Bruker Daltonics) with a nano-electrospray ion source (CaptiveSpray, Bruker Daltonics) in dia-PASEF mode. The dia-PASEF method was defined with a m/z range 400-1201 and a mobility range of 1/K0 = 0.6-1.6 $Vs/cm^2$ using equal ion accumulation and ramp times of 100 ms in the dual TIMS analyser. The collision energy was lowered as a function of increasing ion mobility from 59 eV at 1/K0 = 1.4 $Vs/cm^2$ to 20 eV at 1/K0 = 0.6 $Vs/cm^2$. The diaPASEF window scheme consisted of 32 isolation windows of 26 m/z width (cycle time: 1.8 s) with 1 Da mass overlap and 1 ion mobility window.

### MRT

##### Liquid Chromatography

Peptide samples were loaded onto Evotips using the manufacturers protocol and analysed by the Evosep One LC system (Evosep) using the inbuilt “15 samples per day” method in an identical manner to the Exploris 480 method described above.

##### MRT MSe

The Xevo MRT mass spectrometer was operated in positive electrospray ionisation (ESI) mode. The capillary voltage was 2.8 kV, cone voltage was 30 V and source temperature was set at 100 °C. Data were acquired over m/z range 50-2000 with a scan rate of 1 Hz. All mass spectral data were acquired in continuum mode using MSE to obtain fragmentation data simultaneously[1]. Function one (low energy) data were collected using a constant collision energy of 6 meV whilst the second (high energy) function consisted of a collision energy ramp of 25 to 55 meV. For mass accuracy, leucine enkephalin (m/z = 556.27658) (100 mpg µL-1, 50:50 acetonitrile:water, 0.1 % formic acid) was acquired every 5 min. The lock mass was delivered to the reference sprayer of the MS source using the Waters Reagent Delivery System with a flow rate of 1 µL $min^{-1}$.

#### Spectral Library Generation for Skyline

The spectral library for Waters Vion used in the Skyline[2] analysis was generated from fractionate sample data acquired on the Vion as described above. Data was exported from the Vion instrument in uep format and converted to Waters raw format. The “raw” files were processed through the Waters Protein Lynx Global Server tools as described in the supplementary information producing a “final_fragment.csv” file for each injection. These files were then combined with a bespoke R script, and the resulting table processed through

MAYU[3] from the trans-proteomics-pipeline to estimate and control the false discovery rate. The combined final_fragment.csv file was then imported into Skyline as a spectral library.

#### Vion Data Processed through Skyline

A skyline document was set up with the spectral library generated on the Vion as described above. Vion data for the 1:1, 1:2 and 2:1 ratio samples was imported into Skyline, and an MSstats results table exported.

#### timsTOF; Exploris; ZenoTOF and MRT DIANN processing details

The mass spectrometry proteomics data have been deposited to the ProteomeXchange Consortium via the PRIDE[4] partner repository with the dataset identifier PXD07186961. Briefly for each complete data set of triplicate injections of each ratio 1:1, 1:2 and 2:1 by total peptide mass, of *E. coli* and *H. bluephagenesis* from the timsTOF HT; Exploris 480; ZenoTOF 7600 and SELECT SERIES MRT instruments comprising were processed with DIANN v2.3.0. The MRT data was denoised and centroided using OpenMS tools prior to processing with DIANN, all other data was processed directly from proprietary raw data format. The fasta protein sequence database is included in the data deposition and was a concatenation of the combined reference proteome for *E. coli* K12 (UP000000625) and only available proteome for *H. bluephagenesis* sp. TD01 (UP000838464, now available through EBI UniParc) downloaded from uniprot (https://www.uniprot.org/). DIANN was run with a *q*-value threshold of 1%, a precursor m/z range 400-2000, fragment m/z range 100-2000, precursor charges 2-4, up to 1 missed cleavage, up to two variable modifications of oxidised methionine, constant carbamidomethylation of cysteine, and match between runs enabled. The resulting "report.parquet" files were imported into R, prepared for processing using MSstats with the "DIANNtoMSstatsFormat" function and then processed with MSstats which provided absolute protein estimates per sample on a log10 scale, and ratio estimates between samples. The resulting quantification tables were used to produce the figures reported in the Results and Discussion section in the main text.

### Supplementary Figures

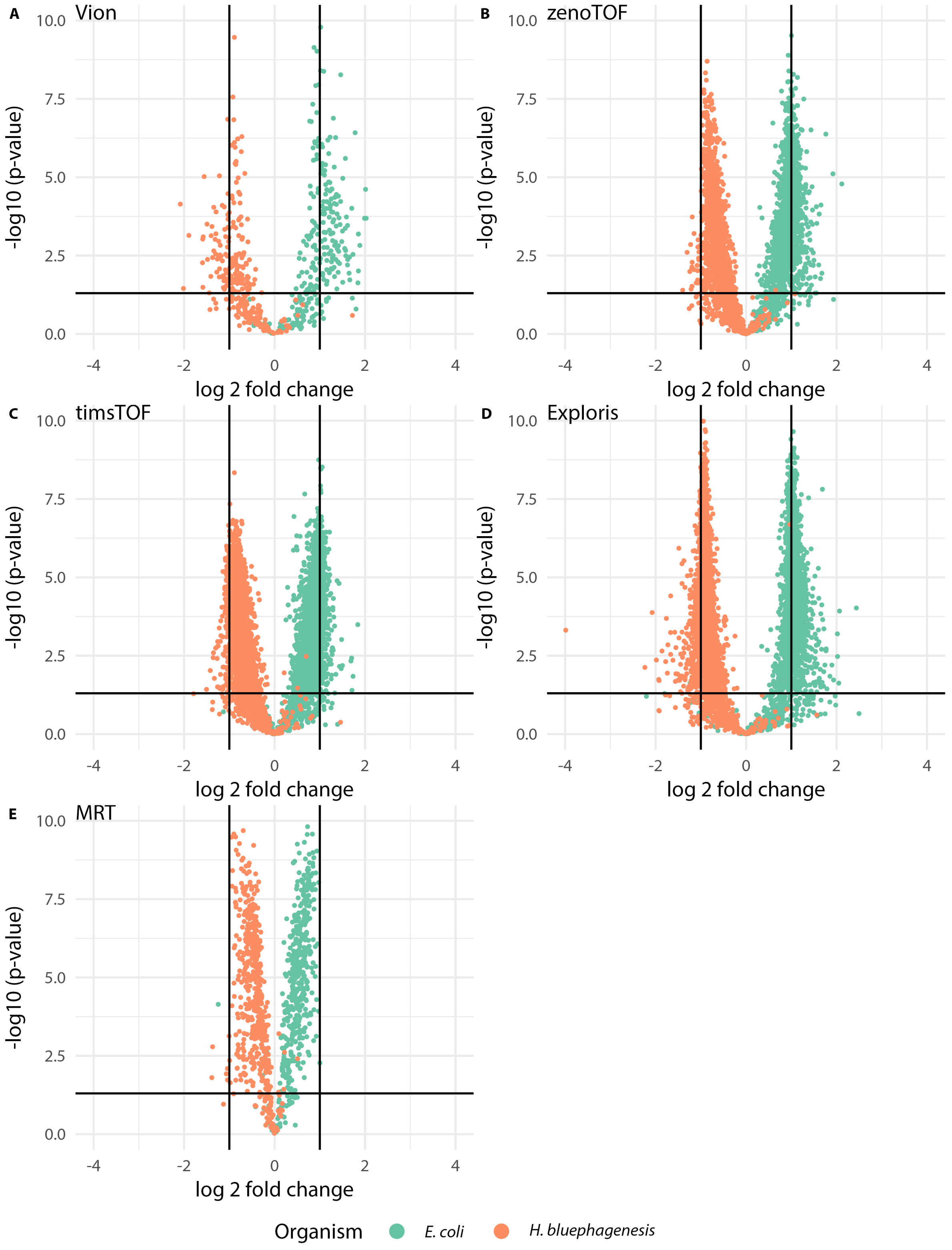


Figure S1: Replot of the volcano plots in Figure 1 with wider x-axis and in larger format to facilitate comparison. Plots data are from A: Vion B: ZenoTOF C: timmsTOF D: Exploris and E: MRT.

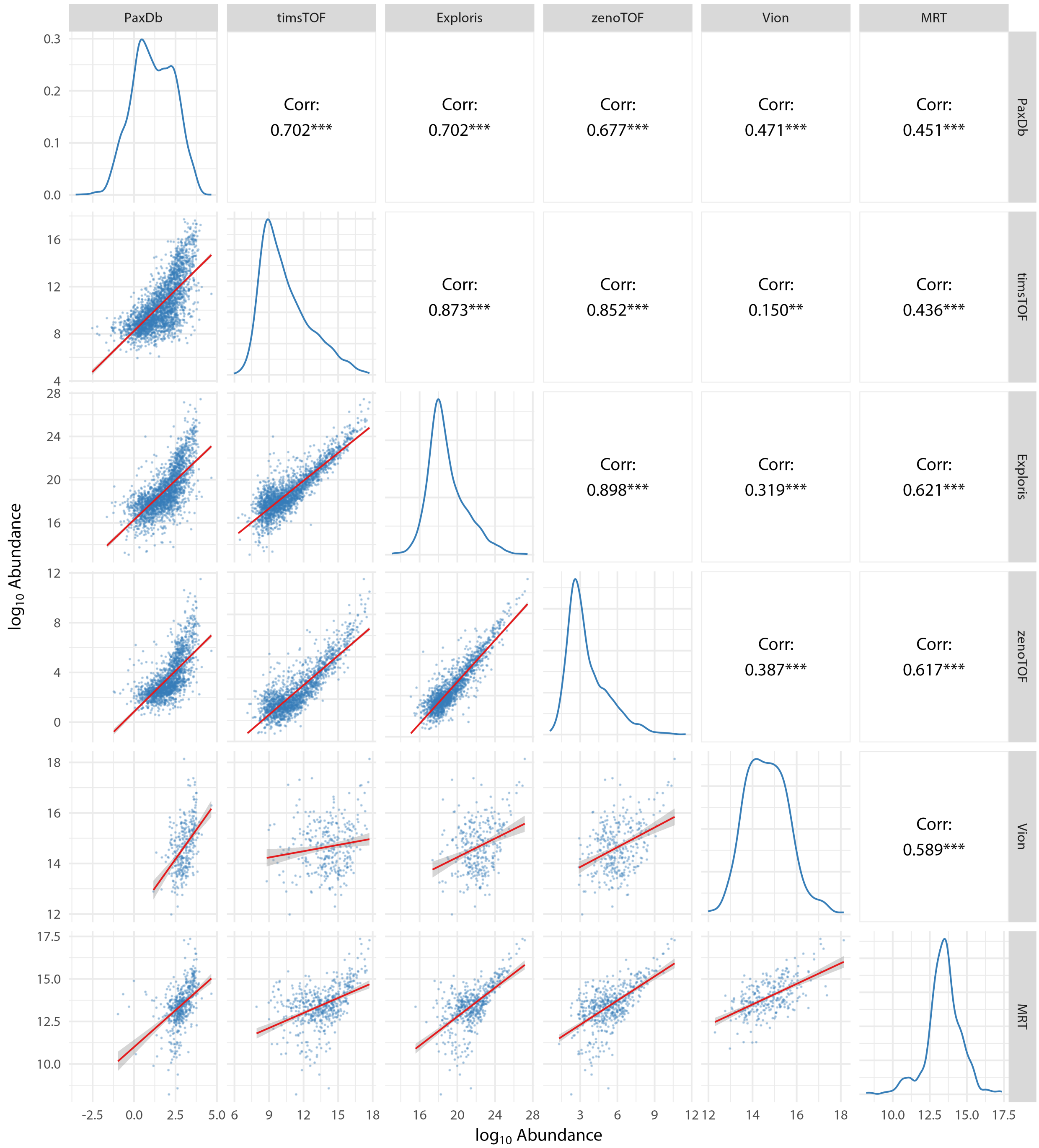


Figure S2: Plot of $\log_{10}$ protein abundance estimates from MSstats analysis of *E. coli* data from each of the instruments and the PaxDb data base. The PaxDb data provides a summary of current estimates of protein expression against which to judge the performance of each instrument. The timsTOF, zenoTOF and Exploris all quantify proteins across the full expression range of PaxDb; the Vion and MRT show they are missing quantification proteins ascribed lower abundance by PaxDb. The plot also enables comparison of instruments with each other, for which the timsTOF, zenoTOF and Exploris give very comparable results, the MRT and Vion showing more deviation, probably due to the absence of lower expression proteins.
