## Supplementary material for "From Discovery Proteomics to Process-Informed Monitoring in Biomanufacturing Chassis Development": Supplimental Information - Documented Scripts


#### Introduction

All the data processing from raw data to figure for this paper[?] was performed with command line tools and R. For the Vion data raw data was processed to a spectral library using the proprietary tools from Waters as ducmented in this paper, the raw data was then read into Skyline against those spectral libraries and the data exported for further analysis by msstats in R. All other data was processed using DIANN v2.3, and the data exported to msstats for further analysis. The script that ran the command lines was written for `GNU Make 4.3`. The full script used to process the data is fully documented below.

#### Set Shell

Each recipe in the make file is run in a "shell", that is a separate command line environment. Here the `make` is run in Ubunto on windows subsystem for linux and the shell is set to `bash`. Those proprietary Specifying the `.ONESHELL` target here with no dependencies instructs `make` to run recipe lines in the same shell instance, rather than each line in a new shell, that is important for a few recipes where one line will specify a variable used in a subsequent line.

```
SHELL = /bin/bash
.ONESHELL:
```

#### Target Output

The final targets are listed at the top of the file. This is only a list, they are specified as the default targets for the `makefile` below.

```
# Path to all mass spec data
allRawData = /mnt/e/massSpecData/25_11__PositioningProteomics/
allOutputs = ${allRawData}

# /mnt/e/massSpecData/25_11__PositioningProteomics/outputLibrary_tpp.splib

exp_blib := ${allOutputs}/outputLibrary_sky.blib
exp_splib := ${allOutputs}/outputLibrary_tpp.splib



allTargets=${exp_blib} ${exp_splib}
```

#### Paths to Executables

Explicit control of software versions is assured by assigning the full path of each executable to a macro for each of the programs required for the pipeline. These must be edited to be correct for the system on which the make script is to be run. Setting them explicitly should also support reproducibility since the version of software used here may be installed along side subsequent releases of the software being used in future active research. Checking these are correct also acts as a check list to ensure system is set up correctly.

```
# paths to the Waters executable, given in windows format
# as passed to windows cmd.exe as a string.

# UEP to MassLynxRaw converter v0.1.4
uep2raw_path=\
"D:/software/UepUdfRawFileConverter-0.1.4/UepUdfConverter/UepUdfToMassLynxRaw.exe"

#ApexRT version: 1.14.0.0 Compiled: Aug  3 2022 12:41:05
apex_path="D:/software/WatersSoftwarePipeline_11_22/ApexRT_1.14.0.0/ApexRT.exe"



# iDTs.exe version 1.79 compiled: 09/06/2022  15:07
idts_path="D:/software/WatersSoftwarePipeline_11_22/iDTs_v1.79/iDTs.exe"

#IADBs.exe version 2.164.16 compiled: 27/06/2022  14:13
IADB_path="D:/software/WatersSoftwarePipeline_11_22/IADBs_v2.164.16/IADBs.exe"

# Waters Csv2Mzid Data Converter, version 0.2 (beta)
```

```
Csv2Mzid="D:/software/Waters_fragcsv2mzid/Csv2Mzid_v0.2Beta/Csv2Mzid.exe"

# Paths to R version 4.3.1 (2023-06-16) -- "Beagle Scouts"
R_path=/usr/bin/R


# Paths to BiblioSpec executables
buildblib_path="D:/software/BiblioSpec/BlibBuild.exe"
buildblibFilter_path="D:/software/BiblioSpec/BlibFilter.exe"
blib2ms2_path="D:/software/BiblioSpec/BlibToMs2.exe"


# Paths to TPP executables
spectrast_path="/usr/local/tpp/bin/spectrast"
Mayu_path="/usr/local/tpp/bin/Mayu.pl"

# Path to diann
diann_path="C:/DIA-NN/2.3.1/diann.exe"
```

#### Macros

In `make` "macros" are rules for converting one text string, or a list of strings to another. For example the macro `watFilePath` takes a list of waters raw data directories such as `myRawDir.raw` and appends the expected contents of the dir `/_header.txt` etc. Listing all intermediate files ensures that if a process does not complete, or a file is deleted or updated, `make` will rebuild the required files. Also having macros defined here enables their re-use in specifying subsequent recipes.

```
timestamp:= $(shell date +%y_%m_%d_%H_%M)
#$(info my timestamp: ${timestamp})

ifdef notestamp
timeStampBackupDir:=./backup/${timestamp}${notestamp}/
define backup
 ${timeStampBackupDir}
endef

endif

define makeDir
$(foreach target,${1},\
if [ ! -d "$(subst /,\,$(abspath $(dir ${target})))"];\
 then mkdir $(subst /,\,$(abspath $(dir ${target}))); fi;\
)
endef



define winify
$(shell readlink -m ${1} | xargs wslpath -m)
endef


UUID=$(shell uuid -v1)
```

```
define watFilePath
$(foreach rawFile,${1},\
$(addprefix ${rawPath}/,$(addsuffix /_header.txt,${rawFile})) \
$(addprefix ${rawPath}/,$(addsuffix /_extern.inf,${rawFile})) \
$(addprefix ${rawPath}/,$(addsuffix /_func001.cdt,${rawFile})) \
$(addprefix ${rawPath}/,$(addsuffix /_func001.dat,${rawFile})) \
$(addprefix ${rawPath}/,$(addsuffix /_func001.idx,${rawFile})) \
$(addprefix ${rawPath}/,$(addsuffix /_func001.ind,${rawFile})) \
$(addprefix ${rawPath}/,$(addsuffix /_func001.sts,${rawFile})) \
$(addprefix ${rawPath}/,$(addsuffix /_func002.cdt,${rawFile})) \
$(addprefix ${rawPath}/,$(addsuffix /_func002.dat,${rawFile})) \
$(addprefix ${rawPath}/,$(addsuffix /_func002.idx,${rawFile})) \
$(addprefix ${rawPath}/,$(addsuffix /_func002.ind,${rawFile})) \
$(addprefix ${rawPath}/,$(addsuffix /_func002.sts,${rawFile})) \
$(addprefix ${rawPath}/,$(addsuffix /_func003.dat,${rawFile})) \
$(addprefix ${rawPath}/,$(addsuffix /_func003.idx,${rawFile})) \
$(addprefix ${rawPath}/,$(addsuffix /_func003.sts,${rawFile})) \
$(addprefix ${rawPath}/,$(addsuffix /_functns.inf,${rawFile})) \
$(addprefix ${rawPath}/,$(addsuffix /drift_alignment_adjustment.csv,${rawFile})) \
$(addprefix ${rawPath}/,$(addsuffix /drift_calibration_adjustment.csv,${rawFile})) \
$(addprefix ${rawPath}/,$(addsuffix /mob_cal.csv,${rawFile})))
endef


define watFileSkyPath
$(foreach rawFile,${1},\
$(addprefix ${rawPath}/${rawSkyPath}/,$(addsuffix /_header.txt,${rawFile})) \
```

```
$(addprefix ${rawPath}/${rawSkyPath}/,$(addsuffix /_extern.inf,${rawFile})) \
$(addprefix ${rawPath}/${rawSkyPath}/,$(addsuffix /_func001.cdt,${rawFile})) \
$(addprefix ${rawPath}/${rawSkyPath}/,$(addsuffix /_func001.dat,${rawFile})) \
$(addprefix ${rawPath}/${rawSkyPath}/,$(addsuffix /_func001.idx,${rawFile})) \
$(addprefix ${rawPath}/${rawSkyPath}/,$(addsuffix /_func001.ind,${rawFile})) \
$(addprefix ${rawPath}/${rawSkyPath}/,$(addsuffix /_func001.sts,${rawFile})) \
$(addprefix ${rawPath}/${rawSkyPath}/,$(addsuffix /_func002.cdt,${rawFile})) \
$(addprefix ${rawPath}/${rawSkyPath}/,$(addsuffix /_func002.dat,${rawFile})) \
$(addprefix ${rawPath}/${rawSkyPath}/,$(addsuffix /_func002.idx,${rawFile})) \
$(addprefix ${rawPath}/${rawSkyPath}/,$(addsuffix /_func002.ind,${rawFile})) \
$(addprefix ${rawPath}/${rawSkyPath}/,$(addsuffix /_func002.sts,${rawFile})) \
$(addprefix ${rawPath}/${rawSkyPath}/,$(addsuffix /_func003.dat,${rawFile})) \
$(addprefix ${rawPath}/${rawSkyPath}/,$(addsuffix /_func003.idx,${rawFile})) \
$(addprefix ${rawPath}/${rawSkyPath}/,$(addsuffix /_func003.sts,${rawFile})) \
$(addprefix ${rawPath}/${rawSkyPath}/,$(addsuffix /_functns.inf,${rawFile})))
endef

define skyRawToOrigRaw
$(foreach rawFile,${1},\
$(subst /${rawSkyPath}/,/,${rawFile}))
endef



#The flowing macro allows .uep files to be stored in sub dir if required.
define watUepFilePath
$(abspath $(addprefix ${allRawData},${1}))
endef
```

```
define apexOut_1
$(foreach raw,${1},\
$(raw:.raw/_header.txt=.raw)/Apex_1/$(notdir $(raw:.raw/_header.txt=_func001.csv)) \
$(raw:.raw/_header.txt=.raw)/Apex_1/$(notdir $(raw:.raw/_header.txt=_func002.csv)) \
)
endef

define idtsOut_1
$(foreach raw,${1},\
$(raw:.raw/_header.txt=.raw)/Apex_1/IDTs_1/$(notdir $(raw:.raw/_header.txt=_iDTs_CPPIS.m
$(raw:.raw/_header.txt=.raw)/Apex_1/IDTS_1/$(notdir $(raw:.raw/_header.txt=_iDTs_CPPIS_S
$(raw:.raw/_header.txt=.raw)/Apex_1/IDTS_1/$(notdir $(raw:.raw/_header.txt=_iDTs_PccChai
$(raw:.raw/_header.txt=.raw)/Apex_1/IDTS_1/$(notdir $(raw:.raw/_header.txt=_iDTs_SSPPIS.
)
endef

define idtsOut_2
$(foreach raw,${1},\
$(raw:.raw/_header.txt=.raw)/Apex_1/IDTs_2/$(notdir $(raw:.raw/_header.txt=_iDTs_CPPIS.m
$(raw:.raw/_header.txt=.raw)/Apex_1/IDTS_2/$(notdir $(raw:.raw/_header.txt=_iDTs_CPPIS_S
$(raw:.raw/_header.txt=.raw)/Apex_1/IDTS_2/$(notdir $(raw:.raw/_header.txt=_iDTs_PccChai
$(raw:.raw/_header.txt=.raw)/Apex_1/IDTS_2/$(notdir $(raw:.raw/_header.txt=_iDTs_SSPPIS.
)
endef
```

```
define iadbOut_1
$(foreach raw,${1},\
$(raw:.raw/_header.txt=.raw)/Apex_1/IDTs_1/$(notdir $(raw:.raw/_header.txt=_iDTs_CPPIS_S
$(raw:.raw/_header.txt=.raw)/Apex_1/IDTs_1/$(notdir $(raw:.raw/_header.txt=_iDTs_CPPIS_S
$(raw:.raw/_header.txt=.raw)/Apex_1/IDTs_1/$(notdir $(raw:.raw/_header.txt=_iDTs_CPPIS_S
$(raw:.raw/_header.txt=.raw)/Apex_1/IDTs_1/$(notdir $(raw:.raw/_header.txt=_iDTs_CPPIS_S
$(raw:.raw/_header.txt=.raw)/Apex_1/IDTs_1/$(notdir $(combinedFasta:.fasta=_subset.fas))
)
endef

define iadbOut_2
$(foreach raw,${1},\
$(raw:.raw/_header.txt=.raw)/Apex_1/IDTS_2/$(notdir $(raw:.raw/_header.txt=_iDTs_CPPIS_S
$(raw:.raw/_header.txt=.raw)/Apex_1/IDTS_2/$(notdir $(raw:.raw/_header.txt=_iDTs_CPPIS_S
$(raw:.raw/_header.txt=.raw)/Apex_1/IDTS_2/$(notdir $(raw:.raw/_header.txt=_iDTs_CPPIS_S
$(raw:.raw/_header.txt=.raw)/Apex_1/IDTS_2/$(notdir $(raw:.raw/_header.txt=_iDTs_CPPIS_S
$(raw:.raw/_header.txt=.raw)/Apex_1/IDTS_2/$(notdir $(combinedFasta:.fasta=_subset.fas))
)
endef



# The following macros turns a space separated list of from make into a quoted comma sep
# It allows lists specified by make to be processed by R scripts.
define RlistIfy
$(shell echo -e "$(foreach string,${1},'${string}',)" | head --bytes=-2)
```

```
endef


# The following macros define empty space and new lines to enable file lists to be writt
# to a text file for import into a second program.


blank :=
define newline


$(blank)
endef


tab := $(shell printf '\011')
```

#### Mass Spectrometry Data Files

Mass spectrometry data are stored as multiple files inside a folder. `Make` can't treat a directory as a pre-requisite so each file is specified as the `_header.txt` file held in each of the directories and this element is removed when passed to waters programs.

The raw data file paths are concatenated into a single list of files so they can each be separately processed by a single unchanging command line. This ensures consistency in data processing.

##### Waters Vion

The raw data from the Vion was exported in "uep" format. That was converted to the Waters "raw" format for processing.

```
# sets the location of the raw data. It can be useful to set this up as a symlink.
rawPath := ${allRawData}
rawSkyPath := rawSky

# The .uep files contain the data exported from Unifi, they must be converted into .rawf
rawUep:= 23_03_06__mbpssmr9_Eco_Halo_Batch7.uep 23_03_06_mbpssmr9_Eco_Halo_Batch6.uep \
        23_03_06__mbpssmr9_Eco_Halo_Batch5.uep 23_03_06__mbpssmr9_Eco_Halo_Batch4.uep \
        23_03_06_mbpssmr9_Eco_Halo_Batch3.uep 23_03_06__mbpssmr9_Eco_Halo_Batch2.uep \
        23_03_06__mbpssmr9_Eco_Halo_Batch1.uep 23_03_06_mbpssmr9_Eco_Halo_Batch8.uep



rawDir_B1 := Inj_1_Ecoli_F1_B1_1.raw Inj_2_Rat_E1_1H_B1_2.raw Inj_3_Ecoli_F1_B1_3.raw \
                Inj_4_Ecoli_F4_B1_4.raw Inj_5_Rat_E1_1H_B1_5.raw Inj_6_Rat_E1_2H_B1_6.ra
                Inj_7_Halo_F1_B1_7.raw Inj_8_Rat_E2_1H_B1_8.raw Inj_9_Halo_F6_B1_9.raw \
                Inj_10_Ecoli_F3_B1_10.raw

rawDir_B2 := Inj_11_Halo_F5_B2_1.raw Inj_12_Halo_F5_B2_2.raw Inj_13_Halo_F2_B2_3.raw \
            Inj_14_Ecoli_F8_B2_4.raw Inj_15_Halo_pure_B2_5.raw Inj_16_Halo_F1_B2_6.raw \
            Inj_17_Ecoli_F4_B2_7.raw Inj_18_Ecoli_F8_B2_8.raw \
            Inj_19_Halo_F8_B2_9.raw Inj_20_Halo_pure_B2_10.raw Inj_21_Halo_F3_B2_11.raw

rawDir_B3 := Inj_22_Ecoli_F2_B3_1.raw Inj_23_Halo_F4_B3_2.raw Inj_24_Halo_F2_B3_3.raw \
            Inj_25_Ecoli_F2_B3_4.raw Inj_26_Ecoli_F7_B3_5.raw Inj_27_Rat_E2_1H_B3_6.raw
            Inj_28_Ecoli_F2_B3_7.raw Inj_29_Halo_F3_B3_8.raw Inj_30_Halo_F7_B3_9.raw \
            Inj_31_Ecoli_F5_B3_10.raw Inj_32_Halo_pure_B3_11.raw Inj_33_Rat_E1_2H_B3_12.
```

```
rawDir_B4 := Inj_34_Halo_F6_B4_1.raw Inj_35_Halo_F1_B4_2.raw Inj_36_Halo_F3_B4_3.raw \
            Inj_37_Halo_F5_B4_4.raw Inj_38_Ecoli_F7_B4_5.raw Inj_39_Ecoli_F3_B4_6.raw \
            Inj_40_Ecoli_F6_B4_7.raw Inj_41_Eco_pure_B4_8.raw Inj_42_Halo_F2_B4_9.raw \
            Inj_43_Rat_E1_2H_B4_10.raw

rawDir_B5 := Inj_43_Rat_E1_2H_B5_1.raw Inj_44_Eco_pure_B5_2.raw Inj_45_Ecoli_F4_B5_3.raw
            Inj_46_Ecoli_F5_B5_4.raw Inj_47_Halo_F6_B5_5.raw Inj_48_Rat_E2_1H_B5_6.raw \
            Inj_49_Halo_F4_B5_7.raw Inj_50_Ecoli_F1_B5_8.raw Inj_51_Ecoli_F7_B5_9.raw \
            Inj_52_Halo_F7_B5_10.raw Inj_53_Halo_F4_B5_11.raw

rawDir_B6 := Inj_54_Ecoli_F8_B6_1.raw Inj_55_Ecoli_F6_B6_2.raw Inj_56_Ecoli_F3_B6_3.raw
            Inj_57_Ecoli_F6_B6_4.raw Inj_58_Halo_F8_B6_5.raw Inj_59_Ecoli_F5_B6_6.raw \
            Inj_60_Halo_F7_B6_7.raw Inj_61_Halo_F8_B6_8.raw Inj_62_Eco_pure_B6_9.raw \
            Inj_63_Rat_E1_1H_B6_10.raw

rawDir_B7 := Inj_64_Rat_E2_1H_B7_1.raw Inj_65_Rat_E1_1H_B7_2.raw Inj_66_Rat_E1_2H_B7_3.r
                Inj_67_Eco_pure_B7_4.raw Inj_68_Halo_pure_B7_5.raw Inj_69_Rat_E2_1H_B7_6
                Inj_70_Rat_E1_1H_B7_7.raw Inj_71_Rat_E1_2H_B7_8.raw Inj_72_Eco_pure_B7_9
                Inj_73_Halo_pure_B7_10.raw

rawDir_B8 := Inj_74_Rat_E2_1H_B8_1.raw Inj_75_Rat_E1_1H_B8_2.raw Inj_76_Rat_E1_2H_B8_3.r
            Inj_77_Eco_pure_B8_4.raw Inj_78_Halo_pure_B8_5.raw Inj_79_Rat_E2_1H_B8_6.raw
            Inj_80_Rat_E1_1H_B8_7.raw Inj_81_Rat_E1_2H_B8_8.raw Inj_82_Eco_pure_B8_9.raw
             Inj_83_Halo_pure_B8_10.raw



rawDir := ${rawDir_B1} ${rawDir_B2} ${rawDir_B3} ${rawDir_B4} \
```

```
            ${rawDir_B5} ${rawDir_B6} ${rawDir_B7} ${rawDir_B8}

rawHead := $(addprefix ${rawPath}/,$(addsuffix /_header.txt,${rawDir}))

FILTER = $(foreach v,$(2),$(if $(findstring $(1),$(v)),$(v),))

rawForSky := $(call FILTER,pure,${rawDir}) $(call FILTER,Rat,${rawDir})

rawFilesSkyline := $(call watFileSkyPath,${rawForSky})
```

#### Waters MRT

Data was exported from the MRT instrument directly as "mzML" files.

```
# sets the location of the raw data. It can be useful to set this up as a symlink.
# ln -s </target-directory> </symlink-directory/example-symlink>
MRT_RawPath :=  ${allRawData}

MRT_RawAll:=  UNIMAN_30SPD_400ng_009_cl_pw_noZ_pw_noZ_reComv.mzML UNIMAN_30SPD_400ng_007
            UNIMAN_30SPD_400ng_008_cl_pw_noZ_pw_noZ_reComv.mzML UNIMAN_30SPD_400ng_006_c
            UNIMAN_30SPD_400ng_005_cl_pw_noZ_pw_noZ_reComv.mzML UNIMAN_30SPD_400ng_001_c
            UNIMAN_30SPD_400ng_004_cl_pw_noZ_pw_noZ_reComv.mzML UNIMAN_30SPD_400ng_002_c
            UNIMAN_30SPD_400ng_003_cl_pw_noZ_pw_noZ_reComv.mzML

MRT_Raw := $(addprefix ${MRT_RawPath}/,${MRT_RawAll})
```

##### Sciex Zenotof Data

Sciex data is prcessed directly from the “wiff” format.

```
# sets the location of the raw data. It can be useful to set this up as a symlink.
# ln -s </target-directory> </symlink-directory/example-symlink>
wiffPath :=  ${allRawData}

wiffAll:= 01_SWATH_2E_1H.wiff 02_SWATH_1E_1H.timeseries.data 02_SWATH_1E_1H.wiff \
        03_SWATH_1E_2H.wiff 04_SWATH_2E_1H.wiff 05_SWATH_1E_1H.wiff 06_SWATH_1E_2H.wiff
        07_SWATH_2E_1H.wiff 08_SWATH_1E_1H.wiff 09_SWATH_1E_2H.wiff

wiffPath := $(addprefix ${wiffPath}/,${wiffAll})

wiff:=$(filter-out %2,$(filter %.wiff,${wiffPath}))
```

##### Thermo Exploris

Thermo data is processed directly from the “raw” format.

```
# sets the location of the raw data. It can be useful to set this up as a symlink.
# ln -s </target-directory> </symlink-directory/example-symlink>
ThermoRawPath :=  ${allRawData}

ThermoRawAll := PB_051223_1E2H_1.raw PB_051223_1E2H_2.raw PB_051223_1E2H_3.raw \
                PB_051223_1to1_1.raw PB_051223_1to1_2.raw PB_051223_1to1_3.raw \
                PB_051223_2E1H_1.raw PB_051223_2E1H_2.raw PB_051223_2E1H_3.raw
```

```
ThermoRaw := $(addprefix ${ThermoRawPath}/,${ThermoRawAll})
```

##### timsTOF Data

The timsTOF data is processed directly fromt the "d" data format.

```
# sets the location of the raw data. It can be useful to set this up as a symlink.
# ln -s </target-directory> </symlink-directory/example-symlink>
Timms_d_path := ${allRawData}



Timms_dAll := PB_051223_1E2H_1_Slot1-19_1_2143.d  PB_051223_1E2H_3_Slot1-19_1_2149.d  PB
                            PB_051223_2E1H_1_Slot1-21_1_2145.d  PB_051223_2E1H_3_Slot1-2
                            PB_051223_1to1_1_Slot1-20_1_2144.d  PB_051223_1to1_3_Slot1-2



Timms_d := $(addprefix ${Timms_d_path}/,${Timms_dAll})
#$(info Timms_d: ${Timms_d})
```

#### Fasta Files

The `.fasta` protein database is specified here. This comprises the *E. coli* proteome with accession no UP000000625 and the *H. bluephagenesis* proteome with accession no UP000838464 database from Uniprot the iRT peptides and common contaminates from the maxQuant contaminant database.

```
combinedFasta := ./ini/UniParc_UP000838464_uniprotkb_proteome_UP000000625_2025_11_10_cRA
combinedDIANNsplib := ./ini/UniParc_UP000838464_uniprotkb_proteome_UP000000625_2025_11_1
combinedFastaRand := $(combinedFasta:.fasta=Rand.fasta)
combinedFastaRandAnnotate := $(combinedFasta:.fasta=Rand_Annot.fasta)
```

#### Intermediate Files

Lists of intermediate files are required for to drive recipes for files along the pathway. Here the names of the required intermediate files are determined from the raw data file names using the macros above.

The following lists the intermediate friles required for processing the Vion data.

```
# The list of files produced by APEX
apex_1 := $(call apexOut_1,${rawHead})
${apex_1}
# The list of files produced by iDTs, in two variants of the processing
idts_1 := $(call idtsOut_1,${rawHead})
idts_2 := $(call idtsOut_2,${rawHead})



# The iadb search results from each of the iDTs described above.
iadb_1 := $(call iadbOut_1,${rawHead})
iadb_2 := $(call iadbOut_2,${rawHead})



subsetFasta=$(filter %_subset.fas, ${iadb_1} ${iadb_2})
```

```
#$(info subsetFasta: ${subsetFasta})

# The names of the combined final fragment files,
# both the .csv and a binary version for R.
iadb_all_final_frag := $(filter %_IA_final_fragment.csv,${iadb_1} ${iadb_2})
iadb_all_final_frag_R := $(call RlistIfy,${iadb_all_final_frag})



# The names of the final frag mzid
iadb_all_final_frag_mzid:=$(iadb_all_final_frag:.csv=.mzid)



# The files produced by Mayu
Mayu_csv := ${allRawData}/Mayu.csv
Mayu_out_txt := ${allRawData}/mayuOutput.txt
combinedMayu_psm := ${allRawData}/combinedMayu_psm_mFDR0.1_t_1.07.csv

final_frac_csv := ${allRawData}/compsite_library_final_fragment.csv
final_frac_rds := ${allRawData}/compsite_library_final_fragment.rds
final_frac_mzid := ${allRawData}/compsite_library_final_fragment.mzid

tables := ${Mayu_csv} ${final_frac_csv}
rds := ${final_frac_rds}
```

### DIA-NN Output Files

The following lists the output files for prcoessing by DIANN.

```
# The zenoTOF and DIANN related files


diannBaseRep_zeno := ./diann_out2/zeno/report_zeno

diannBaseRep_timms := ./diann_out2/timms/report_timms

diannBaseRep_exploris := ./diann_out2/exploris/report_exploris

diannBaseRep_mrt := ./diann_out2/mrt/report_mrt


diannLibs:=  ${combinedDIANNsplib} ${combinedDIANNtsv} ${combinedDIANNparquetLib}


diann_zenoTOF := \
    $(addsuffix .gg_matrix.tsv,${diannBaseRep_zeno}) \
    $(addsuffix .log.txt,${diannBaseRep_zeno}) \
    $(addsuffix -qn.parquet,${diannBaseRep_zeno}) \
    $(addsuffix .pr_matrix.tsv,${diannBaseRep_zeno}) \
    $(addsuffix .pg_matrix.tsv,${diannBaseRep_zeno}) \
    $(addsuffix .protein_description.tsv,${diannBaseRep_zeno}) \
    $(addsuffix -lib.parquet,${diannBaseRep_zeno})


diann_ThermoRaw := \
    $(addsuffix .gg_matrix.tsv,${diannBaseRep_exploris}) \
    $(addsuffix .log.txt,${diannBaseRep_exploris}) \
    $(addsuffix -qn.parquet,${diannBaseRep_exploris}) \
    $(addsuffix .pr_matrix.tsv,${diannBaseRep_exploris}) \
    $(addsuffix .pg_matrix.tsv,${diannBaseRep_exploris}) \
```

```
    $(addsuffix .protein_description.tsv,${diannBaseRep_exploris}) \
    $(addsuffix -lib.parquet,${diannBaseRep_exploris})

diann_Timms_d := \
    $(addsuffix .gg_matrix.tsv,${diannBaseRep_timms}) \
    $(addsuffix .log.txt,${diannBaseRep_timms}) \
    $(addsuffix -qn.parquet,${diannBaseRep_timms}) \
    $(addsuffix .pr_matrix.tsv,${diannBaseRep_timms}) \
    $(addsuffix .pg_matrix.tsv,${diannBaseRep_timms}) \
    $(addsuffix .protein_description.tsv,${diannBaseRep_timms}) \
    $(addsuffix -lib.parquet,${diannBaseRep_timms})

diann_MRT := \
    $(addsuffix .gg_matrix.tsv,${diannBaseRep_mrt}) \
    $(addsuffix .log.txt,${diannBaseRep_mrt}) \
    $(addsuffix -qn.parquet,${diannBaseRep_mrt}) \
    $(addsuffix .pr_matrix.tsv,${diannBaseRep_mrt}) \
    $(addsuffix .pg_matrix.tsv,${diannBaseRep_mrt}) \
    $(addsuffix .protein_description.tsv,${diannBaseRep_mrt}) \
    $(addsuffix -lib.parquet,${diannBaseRep_mrt}) \



wiff_dia:=$(addprefix ./dia/,$(notdir $(wiff:.wiff=.wiff.dia)))
Timms_d_dia:=$(addprefix ./dia/,$(notdir $(Timms_d:.d=.d.dia)))
ThermoRaw_dia:=$(addprefix ./dia/,$(notdir $(ThermoRaw:.raw=.raw.dia)))
```

```
diann_all := ${diann_zenoTOF} ${diann_ThermoRaw} \
                ${diann_Timms_d} ${diann_MRT}
```

##### Specify Default Targets

The default target of the `makefile` is the target of the first defined dependency. Here the prerequisites of the `.PHONY` target will be recompiled unconditionally regardless of the existance of a file of tat name or it last-modification time. Here the two targets of `.PHONY` are`all` and `clean`. The `all` target is a list of the final requirements of the makefile. The `clean` target contains rules to remove unwanted side effect files. It is possible to remove all intermediate files but often these are of interest to reivew the process.

```
.PHONY: all clean sky diann_out_target justDiann

all: ${diannLibs} ${allDia} ${diann_all}

justDiann: ${diann_all}
iadbAll : ${iadb_all_final_frag} ${Mayu_out_txt} ${iadb_all_final_frag} ${final_frac_csv
skyAll : ${exp_blib} ${exp_splib} ${rawFilesSkyline}

clean :
    rm -f ${apex_1} ${idts_1} ${idts_2} ${iadb_1} ${iadb_2} ${iadb_all_final_frag} \
    ${iadb_all_final_frag_R} ${iadb_all_final_frag_mzid} \
    ${Mayu_csv} ${Mayu_out_txt} ${combinedMayu_psm} \
    ${final_frac_csv} ${final_frac_rds} ${final_frac_mzid} \
    ${tables} ${rds} \
```

```
    ${diann_zenoTOF} ${diann_ThermoRaw} ${diann_Timms_d} ${diann_MRT} \
    ${wiff_dia} ${Timms_d_dia} ${ThermoRaw_dia} \
    ${exp_blib} ${exp_splib} ${rawFilesSkyline}
```

#### Waters Vion Pipeline

##### Raw Data Converstion

Raw data is exported from the Vion acquisition software "unifi" in a .uep format. That format is converted into Waters `.raw` format for further processing.

```
$(call watFilePath,${rawDir_B1}) &: $(call watUepFilePath,23_03_06__mbpssmr9_Eco_Halo_Ba
    if [ ! -d "./raw" ]; then mkdir -p ${allRawData}; fi
    cmd.exe /c ${uep2raw_path} -a -u $(call winify,$<)

$(call watFilePath,${rawDir_B2}) &: $(call watUepFilePath,23_03_06__mbpssmr9_Eco_Halo_Ba
    if [ ! -d "./raw" ]; then mkdir -p ${allRawData}; fi
    cmd.exe /c ${uep2raw_path} -a -u $(call winify,$<)

$(call watFilePath,${rawDir_B3}) &: $(call watUepFilePath,23_03_06__mbpssmr9_Eco_Halo_Ba
    if [ ! -d "./raw" ]; then mkdir -p ${allRawData}; fi
    cmd.exe /c ${uep2raw_path} -a -u $(call winify,$<)
```

```
$(call watFilePath,${rawDir_B4}) &: $(call watUepFilePath,23_03_06__mbpssmr9_Eco_Halo_Ba
    if [ ! -d "./raw" ]; then mkdir -p ${allRawData}; fi
    cmd.exe /c ${uep2raw_path} -a -u $(call winify,$<)



$(call watFilePath,${rawDir_B5}) &: $(call watUepFilePath,23_03_06__mbpssmr9_Eco_Halo_Ba
    if [ ! -d "./raw" ]; then mkdir -p  ${allRawData}; fi
    cmd.exe /c ${uep2raw_path} -a -u $(call winify,$<)



$(call watFilePath,${rawDir_B6}) &: $(call watUepFilePath,23_03_06__mbpssmr9_Eco_Halo_Ba
    if [ ! -d "./raw" ]; then mkdir -p  ${allRawData}; fi
    cmd.exe /c ${uep2raw_path} -a -u $(call winify,$<)


$(call watFilePath,${rawDir_B7}) &: $(call watUepFilePath,23_03_06__mbpssmr9_Eco_Halo_Ba
    if [ ! -d "./raw" ]; then mkdir -p  ${allRawData}; fi
    cmd.exe /c ${uep2raw_path} -a -u $(call winify,$<)


$(call watFilePath,${rawDir_B8}) &: $(call watUepFilePath,23_03_06__mbpssmr9_Eco_Halo_Ba
    if [ ! -d "./raw" ]; then mkdir -p ${allRawData}; fi
    cmd.exe /c ${uep2raw_path} -a -u $(call winify,$<)
```

##### Peak Picking

The Apex3d and peptide3d algorithms between them work to pick peaks from the raw data, de-isotope them, relate precurser and product ions and produce spectra peaklists for database searching.

##### Apex Feature Picking

The Apex algorithm is run with settings closely mirroring those run by PLGS. The major difference is substantial effort put into optimising the low-energy and high-energy thresholds that appear critical in getting high numbers of peptides hits.

```
define rawApex_rule
${1} &: ${2}
    cmd.exe /c ${apex_path} \
        -pRawDirName "$(call winify,$(2:.raw/_header.txt=.raw))" \
        -outputDirName "$(call winify,$(2:.raw/_header.txt=.raw))/Apex_1" \
        -startingRTMin 5 \
        -endingRTMin 75 \
        -lockMassZ1 556.2771 \
        -lockMassToleranceAMU 1.25 \
        -lmThresholdCounts 500 \
        -msResolution Real-time \
        -chromFWHMMin 0.5 \
        -driftFWHMStart Real-time \
        -driftFWHMEnd Real-time \
        -adcAveResponse 1 \
        -leScanAvgIntensityFactor 0.2 \
        -heScanAvgIntensityFactor 0.1 \
        -leDeltaMz 50 \
        -heDeltaMz 60 \
        -leDeltaDrift 3 \
        -heDeltaDrift 4 \
        -deltaScan 6 \
```

```
        -coeffVariationThreshold 0.1 \
        -dreDependentThreshold 1 \
        -UW 1 \
        -noiseFloorFilter 1 \
        -noiseFloorFilterSNR 3 \
        -enableDiagnostics \
        -showExcludedPeaks 0 \
        -writeSummary
endef



$(foreach raw, ${rawHead},$(eval $(call rawApex_rule,\
$(call apexOut_1,${raw}),\
${raw})))
```

##### Peak Aggregation to Spectra

The feature lists produced by Apex must be agregated into spectra in order to be searched. The iDTs program does this by matching elution profiles and drift times of precurser and product ions.

The algorithm is controled by extensive options outlined in a `.xml` file, see below.

```
define iDTs_control_1
<?xml version="1.0" encoding="UTF-8"?>
<IDTS_RESEARCH_PARAMETERS VERSION="1.76" TITLE="ION-DIRECTED TARGETED SEARCH">
  <SAMPLE_TYPE VALUE="LM"/>
  <USE_SMART_PARAMETER_RULES VALUE="0"/>
  <FORCE_MS1_PROCESSING VALUE="0"/>
```

```
<PROCESS_WITHOUT_ION_MOBILITY_SEPARATION VALUE="0"/>
<MAX_CPUS VALUE="-1"/><!-- "-1" uses all available CPUs -->
<MIN_RET_TIME_MIN VALUE=""/>
<MAX_RET_TIME_MIN VALUE=""/>
<FUNC_1_MIN_MZ VALUE=""/>
<FUNC_1_MAX_MZ VALUE=""/>
<ENABLE_CCSA VALUE="0"/>
<MASS_SEARCH_ERROR_TOL_PPM VALUE="55.9"/>
<DRIFT_SEARCH_ERROR_TOL_BINS VALUE="5"/>
<ION_DETECTIONS_NOISE_FLOOR_FILTER VALUE="1">
  <BIN_CUTOFF VALUE="3"/>
  <SNR_THRESHOLD VALUE="3.21"/>
</ION_DETECTIONS_NOISE_FLOOR_FILTER>
<MS_INTENSITY_AUTO_THRESHOLDING ALGORITHM="TOP_N_IONS">
  <PRECURSOR_ISOTOPE>
      <THRESHOLD_FACTOR VALUE="0.50"/>
  </PRECURSOR_ISOTOPE>
  <TOP_LOG_INTENSITY>
    <LOG10_INTENSITY_RANGE VALUE="4.0"/>
  </TOP_LOG_INTENSITY>
  <TOP_N_IONS>
    <TOP_N VALUE="2500"/>
  </TOP_N_IONS>
  <BIN_FRACTION>
    <BIN_FRACTION_RATIO VALUE="0.15"/>
  </BIN_FRACTION>
  <SNR>
```

```
    <SNR_THRESHOLD VALUE="3.0"/>
  </SNR>
  <MS_SWITCHED_MASSES>
  </MS_SWITCHED_MASSES>
  <MS_SWITCHED_MASS_INTENSITY>
    <MS_SWITCHED_MASS_INTENSITY_FACTOR VALUE="0.3333333"/>
  </MS_SWITCHED_MASS_INTENSITY>
  <MIN_INTENSITY_THRESHOLD VALUE=""/>
</MS_INTENSITY_AUTO_THRESHOLDING>
<MSE_INTENSITY_AUTO_THRESHOLDING ALGORITHM="TOP_N_IONS">
  <APPLY_BY_PRECURSOR_DRIFT VALUE="0"/>
  <PRECURSOR_ISOTOPE>
      <THRESHOLD_FACTOR VALUE="0.025"/>
  </PRECURSOR_ISOTOPE>
  <TOP_LOG_INTENSITY>
    <LOG10_INTENSITY_RANGE VALUE="3.06"/>
  </TOP_LOG_INTENSITY>
  <TOP_N_IONS>
    <TOP_N VALUE="16800"/>
  </TOP_N_IONS>
  <BIN_FRACTION>
    <BIN_FRACTION_RATIO VALUE="0.15"/>
  </BIN_FRACTION>
  <SNR>
    <SNR_THRESHOLD VALUE="3.67"/>
  </SNR>
  <MIN_INTENSITY_THRESHOLD VALUE=""/>
```

```
</MSE_INTENSITY_AUTO_THRESHOLDING>
<MSMS_INTENSITY_AUTO_THRESHOLDING ALGORITHM="TOP_N_IONS">
  <PRECURSOR_ISOTOPE>
      <THRESHOLD_FACTOR VALUE="0.025"/>
  </PRECURSOR_ISOTOPE>
  <TOP_LOG_INTENSITY>
    <LOG10_INTENSITY_RANGE VALUE="3.0"/>
  </TOP_LOG_INTENSITY>
  <TOP_N_IONS>
    <TOP_N VALUE="1500"/>
  </TOP_N_IONS>
  <BIN_FRACTION>
    <BIN_FRACTION_RATIO VALUE="0.15"/>
  </BIN_FRACTION>
  <SNR>
    <SNR_THRESHOLD VALUE="3.0"/>
  </SNR>
  <MIN_INTENSITY_THRESHOLD VALUE=""/>
</MSMS_INTENSITY_AUTO_THRESHOLDING>
<QUAD_MS_MS_TRANSMISSION>
  <LOW_MASS_ISOLATION_WINDOW VALUE=""/>
  <HIGH_MASS_ISOLATION_WINDOW VALUE=""/>
  <JOIN_ONLY_PCC_MATCHED_TO_SWITCHED_MASS VALUE="1"/>
  <SWITCHED_MASS_MATCH_TOL_PPM VALUE="100"/>
  <SWITCHED_MASS_CAL_OFFSET_PPM VALUE=""/>
</QUAD_MS_MS_TRANSMISSION>
<HYBRID_SONAR>
```

```
    <CALIBRATION_MODE VALUE="Auto"/>
    <MANUAL_CAL>
      <DRIFT_OFFSET VALUE="0"/>
      <DRIFT_SLOPE VALUE="0"/>
    </MANUAL_CAL>
  </HYBRID_SONAR>
  <PRECURSOR_DEISOTOPING>
    <MIN_Z VALUE="2"/>
    <MAX_Z VALUE="6"/>
    <MIN_NUM_ISOTOPES VALUE=""/>
    <MZ_FWHM_WINDOW_FACTOR VALUE="0.5"/>
    <ISOTOPE_INTENSITY_RANGE_FACTOR VALUE="0.3333333"/>
    <MAKE_VIRTUAL_IONS VALUE="1"/>
    <DEISOTOPE_WITH_DRIFT>
      <INTENSITY_SAT_LOG10_RANGE VALUE="1.5"/>
      <MASS_DRIFT_TO_CHARGE_FRAC_MASS_TOL VALUE="40"/><!-- mDa -->
      <CHARGE_1_MAX_MZ_DRIFT_RATIO VALUE="8.8"/>
      <RING_DRIFT_FWHM_WINDOW_FACTOR VALUE="0.25"/>
      <RING_MZ_FWHM_WINDOW_FACTOR VALUE="1.333333"/>
      <RING_INTENSITY_HIGH_TIER_CEILING_FACTOR VALUE="0.50"/>
      <RING_INTENSITY_LOW_TIER_CEILING_FACTOR VALUE="0.15"/>
      <!-- Product Drift offset search range -->
      <PRODUCT_DRIFT_LOW_OFFSET VALUE="-4.5"/>
      <PRODUCT_DRIFT_HIGH_OFFSET VALUE="+0.5"/>
    </DEISOTOPE_WITH_DRIFT>
  </PRECURSOR_DEISOTOPING>
  <PRECURSOR_PEAK_TRACKING>
```

```xml
    <NO_PEAK_REJECTION_FOR_DDA_SPECTRA VALUE="1"/>
    <ALIGN_MS1_INSOURCE_FRAGS_BY_PEAK_RET_TIME VALUE="0"/>
    <PROCESS_ONLY_TOP_N_INTENSE_PRECURSORS VALUE=""/>
    <EXTRACT_PRECURSORS_ABOVE_SNR VALUE="0">
      <SNR_THRESHOLD VALUE="0.59"/>
      <MIN_INTENSITY_CV_FOR_EXTRACTION VALUE="0.25"/>
    </EXTRACT_PRECURSORS_ABOVE_SNR>
    <PEAK_MIN_NUM_SCANS VALUE="3"/>
    <NUM_DROP_OUT_SCANS_TO_TERMINATE_PEAK_CHAIN VALUE="3"/>
    <MIN_PEAK_INTENSITY VALUE="16"/>
    <RATIO_OF_CHANGE_DECIMATION_FACTOR VALUE="2"/>
    <WRITE_PRECURSOR_PEAK_CHAINS_TO_CSV_FILE VALUE="1"/>
    <EXTRACT_PEAKS_WITHOUT_PEAK_PROPERTIES VALUE="1">
      <NUM_SCANS_FOR_PEAK_EXTRACTION_EACH_SIDE_OF_PEAK VALUE="3"/>
    </EXTRACT_PEAKS_WITHOUT_PEAK_PROPERTIES>
    <MIN_NUM_SCANS_FOR_LOW_HIGH_SMR_SELECTION VALUE="5"/>
    <SIGNAL_CV_THRESHOLD_FOR_LOW_HIGH_SMR_SELECTION VALUE="0.20"/>
    <LOW_CV_PEAK_SIGNAL_MEDIAN_RATIO_THRESHOLD VALUE="2.0"/>
    <HIGH_CV_PEAK_SIGNAL_MEDIAN_RATIO_THRESHOLD VALUE="1.0"/>
    <PEAK_MIN_NUM_SCANS_TO_REQUIRE_BOTH_HALF_HEIGHTS VALUE="21"/>
    <PEAK_BASE_WIDTH_EXTRACTION_IN_PLUS_MINUS_STD_DEVS VALUE="3.0"/>
    <ALLOW_PEAKS_WITH_NO_HALF_HEIGHTS VALUE="1"/>
    <MIN_PEAK_LEN VALUE="4"/>
    <MAX_PEAK_LEN VALUE="23"/>
  </PRECURSOR_PEAK_TRACKING>
  <ISOBARIC_LABEL_TRACKING>
    <TRACK_TMT_6PLEX_LABELS VALUE="0">
```

```
    <TMT_MASS_SEARCH_TOL_FACTOR VALUE="2.0"/>
  </TRACK_TMT_6PLEX_LABELS>
</ISOBARIC_LABEL_TRACKING>
<DRIFT_TRANSMISSION_ALIGNMENT VALUE="0">
  <!-- Determines coeluting precursors within drift (mass)
    transmission window for HDMSe/SONAR acquisitions
    and determines relative intensity ratio of each
    precursor then applies that ratio to their respective
    product ion spectrum as a ceiling threshold, extracting
    product ions within specified log intensity range. -->
  <DRIFT_TRANSMISSION_WINDOW_BINS VALUE="2.48"/>
  <PRODUCT_ION_TXM_INTENSITY_FACTOR VALUE="2.87"/>
  <PRODUCT_ION_LOG10_INTENSITY_RANGE VALUE="2.97"/>
</DRIFT_TRANSMISSION_ALIGNMENT>
<PRODUCT_ION_ALIGNMENT>
  <MIN_MZ VALUE="50"/>
  <REMOVE_RESIDUALS_FROM_PRODUCT_ION_SPECTRA VALUE="1"/>
  <REJECT_IONS_INTENSITY_GREATER_THAN_PRECURSOR VALUE="0"/>
  <FIX_SPLIT_IONS VALUE="1"/>
  <REMOVE_ISOTOPES VALUE="1"/>
  <CHARGE_REDUCE_PRODUCT_IONS_TO_MHPLUS VALUE="0"/>
  <ALIGN_PRODUCT_IONS_BY_INTENSITY_RATIO_OF_CHANGE VALUE="0">
    <PRECURSOR_TARGET_ROC VALUE="0.50"/>
    <PRECURSOR_PRODUCT_ION_ROC_MATCHING_FACTOR VALUE="0.40"/>
    <NUM_SCANS_BELOW_HALF_HEIGHTS_TO_STOP_ROC_COMPARES VALUE=""/>
    <TAKE_TOP_N_ROC_SCANS_CLOSEST_TO_TARGET VALUE=""/>
    <REJECT_PRODUCT_IONS_BY_ROC VALUE="0"/>
```

```
    </ALIGN_PRODUCT_IONS_BY_INTENSITY_RATIO_OF_CHANGE>
    <ALIGN_PRODUCT_IONS_BY_MASS_DEFECT VALUE="1">
      <B_ION_HIGH_STRIPE_FACTOR VALUE="1.25"/>
      <Y_ION_LOW_STRIPE_FACTOR VALUE="1.25"/>
    </ALIGN_PRODUCT_IONS_BY_MASS_DEFECT>
    <ALIGN_PRODUCT_IONS_BY_INTENSITY_MASS_BINS VALUE="0">
      <MASS_BIN_WIDTH_SET_BY_MR_DIVIDED_BY_NUM_BINS VALUE="20"/>
      <OVERRIDE_MASS_BIN_WIDTH_AMU VALUE="50"/>
      <KEEP_BIN_IF_NUM_IONS_LESS_THAN_EQ_TO VALUE="2"/>
      <REJECT_BIN_IF_INTENSITY_CV_LESS_THAN VALUE="0.25"/>
      <REJECT_ION_IF_INTENSITY_TO_MEAN_RATIO_LESS_THAN VALUE="0.50"/>
      <TAKE_TOP_N_INTENSE_BIN_IONS VALUE="5"/>
    </ALIGN_PRODUCT_IONS_BY_INTENSITY_MASS_BINS>
  </PRODUCT_ION_ALIGNMENT>
  <CPPIS_ALIGNMENT>
    <!-- MSe SSPPIS Construction Options: only one option can be set -->
    <MAKE_SSPPISS_FROM_MSE_SPECTRA_BETWEEN_HALF_HEIGHTS VALUE="0"/>
    <MAKE_SSPPISS_FROM_ALL_MSE_SPECTRA VALUE="1"/>
    <!-- SSPPIS Filter Options -->
    <USE_SSPPIS_IF_RESIDUAL_FRAG_EFF_GREATER_THAN VALUE=""/>
    <USE_SSPPIS_IF_NO_RESIDUAL_PCC_INTENSITY_GREATER_THAN VALUE=""/>
    <REJECT_SSPPIS_IF_PRECURSOR_INTENSITY_LESS_THAN_RESIDUAL VALUE="0"/><!-- not recomme
    <COMPOSITING_MASS_MATCH_ERROR_TOL_PPM VALUE=""/><!-- defaults to MASS_SEARCH_ERROR_T
    <MIN_NUMBER_OF_SPECTRA_FOR_CPPIS VALUE="3"/>
    <REJECT_COMPOSITE_ION_IF_AR1_CV_GREATER_THAN VALUE=""/><!-- no value disables reject
    <CALC_COMPOSITE_PCC_ISOTOPE_RATIOS_AND_CV VALUE="1"/>
    <ALIGN_PRODUCT_IONS_BY_INTENSITY_MASS_BINS VALUE="1"/><!-- uses PRODUCT_ION_ALIGNMEN
```

```
</CPPIS_ALIGNMENT>

<FILTERED_IONS_FUNC_CSV_FILES_PRODUCTION>

  <WRITE_FILTERED_IONS_FUNC_CSV_FILES VALUE="0"/>

  <WRITE_INCLUDE_EXCLUDE_IONS_AS_SEPARATE_FILES VALUE="1"/>

</FILTERED_IONS_FUNC_CSV_FILES_PRODUCTION>

<SSPPIS_MGF_PRODUCTION>

  <WRITE_SSPPISS_TO_MGF_FILE VALUE="1"/><!-- _iDTs_SSPPISs.mgf -->

  <WRITE_SSPPISS_ONLY_AT_PEAK_HEIGHT VALUE="0"/>

  <WRITE_PCC_ISOTOPE_RATIOS VALUE="1"/>

  <CHARGE_REDUCE_PEPMASS_TO_MHPLUS VALUE="0"/>

  <ADD_PRODUCT_ION_META_DATA VALUE="1"/>

  <SUPPRESS_VIRTUAL_PCC_SPECTRA VALUE="0"/>

  <SUPPRES_PRODUCT_IONS_WITH_ZERO_ROC VALUE="0"/>

  <MIN_NUMBER_PROD_IONS VALUE="3"/>

</SSPPIS_MGF_PRODUCTION>

<CPPIS_MGF_PRODUCTION>

  <WRITE_CPPISS_TO_MGF_FILE VALUE="1"/>

  <WRITE_PCC_ISOTOPE_RATIOS VALUE="1"/>

  <WRITE_PCC_ISOTOPE_RATIOS_CV VALUE="1"/>

  <CHARGE_REDUCE_PEPMASS_TO_MHPLUS VALUE="0"/>

  <ADD_PRODUCT_ION_META_DATA VALUE="1"/>

  <SUPPRESS_VIRTUAL_PCC_SPECTRA VALUE="0"/>

  <MIN_NUMBER_PROD_IONS VALUE="3"/>

</CPPIS_MGF_PRODUCTION>

<PLGS_PRODUCTION>

  <WRITE_CPPISS_TO_SPECTRUM_BIN_FILE VALUE="1"/>

  <COMBINE_CPPISS_INTO_AMRTS VALUE="0"/>
```

```
    <SPLIT_SPECTRUM_BIN_FILE_BY_CHARGE_STATES VALUE="0"/>
    <ADD_CHARGE1_SPECTRA_TO_SPLIT_SPECTRA VALUE="1"/>
    <SUPPRESS_VIRTUAL_PCC_SPECTRA VALUE="0"/>
    <MIN_NUMBER_CPPIS_PROD_IONS VALUE="5"/>
    <WRITE_LE_IDS_CPPIS_IDS_XREF_CSV_FILE VALUE="1"/>
  </PLGS_PRODUCTION>
  <PROGENESIS_PRODUCTION>
    <WRITE_CPPISS_TO_QI_FILES VALUE="0"/>
  </PROGENESIS_PRODUCTION>
</IDTS_RESEARCH_PARAMETERS>
endef

./ini/iDTs_control_1.xml :
    $$(file > ./ini/iDTs_control_1.xml,${iDTs_control_1})
```

```
define iDTs_control_2
<?xml version="1.0" encoding="UTF-8"?>
<IDTS_RESEARCH_PARAMETERS VERSION="1.76" TITLE="ION-DIRECTED TARGETED SEARCH">
  <SAMPLE_TYPE VALUE="LM"/>
  <USE_SMART_PARAMETER_RULES VALUE="0"/>
  <FORCE_MS1_PROCESSING VALUE="0"/>
  <PROCESS_WITHOUT_ION_MOBILITY_SEPARATION VALUE="0"/>
  <MAX_CPUS VALUE="-1"/><!-- "-1" uses all available CPUs -->
  <MIN_RET_TIME_MIN VALUE=""/>
  <MAX_RET_TIME_MIN VALUE=""/>
  <FUNC_1_MIN_MZ VALUE=""/>
  <FUNC_1_MAX_MZ VALUE=""/>
```

```
<ENABLE_CCSA VALUE="0"/>
<MASS_SEARCH_ERROR_TOL_PPM VALUE="60"/>
<DRIFT_SEARCH_ERROR_TOL_BINS VALUE="5"/>
<ION_DETECTIONS_NOISE_FLOOR_FILTER VALUE="1">
  <BIN_CUTOFF VALUE="3"/>
  <SNR_THRESHOLD VALUE="3.0"/>
</ION_DETECTIONS_NOISE_FLOOR_FILTER>
<MS_INTENSITY_AUTO_THRESHOLDING ALGORITHM="TOP_N_IONS">
  <PRECURSOR_ISOTOPE>
      <THRESHOLD_FACTOR VALUE="0.50"/>
  </PRECURSOR_ISOTOPE>
  <TOP_LOG_INTENSITY>
    <LOG10_INTENSITY_RANGE VALUE="4.5"/>
  </TOP_LOG_INTENSITY>
  <TOP_N_IONS>
    <TOP_N VALUE="3000"/>
  </TOP_N_IONS>
  <BIN_FRACTION>
    <BIN_FRACTION_RATIO VALUE="0.15"/>
  </BIN_FRACTION>
  <SNR>
    <SNR_THRESHOLD VALUE="3.0"/>
  </SNR>
  <MS_SWITCHED_MASSES>
  </MS_SWITCHED_MASSES>
  <MS_SWITCHED_MASS_INTENSITY>
    <MS_SWITCHED_MASS_INTENSITY_FACTOR VALUE="0.3333333"/>
```

```
    </MS_SWITCHED_MASS_INTENSITY>
    <MIN_INTENSITY_THRESHOLD VALUE=""/>
  </MS_INTENSITY_AUTO_THRESHOLDING>
  <MSE_INTENSITY_AUTO_THRESHOLDING ALGORITHM="TOP_N_IONS">
    <APPLY_BY_PRECURSOR_DRIFT VALUE="1"/>
    <PRECURSOR_ISOTOPE>
        <THRESHOLD_FACTOR VALUE="0.025"/>
    </PRECURSOR_ISOTOPE>
    <TOP_LOG_INTENSITY>
      <LOG10_INTENSITY_RANGE VALUE="4"/>
    </TOP_LOG_INTENSITY>
    <TOP_N_IONS>
      <TOP_N VALUE="20000"/>
    </TOP_N_IONS>
    <BIN_FRACTION>
      <BIN_FRACTION_RATIO VALUE="0.15"/>
    </BIN_FRACTION>
    <SNR>
      <SNR_THRESHOLD VALUE="3.0"/>
    </SNR>
    <MIN_INTENSITY_THRESHOLD VALUE=""/>
  </MSE_INTENSITY_AUTO_THRESHOLDING>
  <MSMS_INTENSITY_AUTO_THRESHOLDING ALGORITHM="TOP_N_IONS">
    <PRECURSOR_ISOTOPE>
        <THRESHOLD_FACTOR VALUE="0.025"/>
    </PRECURSOR_ISOTOPE>
    <TOP_LOG_INTENSITY>
```

```
      <LOG10_INTENSITY_RANGE VALUE="3.0"/>
    </TOP_LOG_INTENSITY>
    <TOP_N_IONS>
      <TOP_N VALUE="1500"/>
    </TOP_N_IONS>
    <BIN_FRACTION>
      <BIN_FRACTION_RATIO VALUE="0.15"/>
    </BIN_FRACTION>
    <SNR>
      <SNR_THRESHOLD VALUE="3.0"/>
    </SNR>
    <MIN_INTENSITY_THRESHOLD VALUE=""/>
  </MSMS_INTENSITY_AUTO_THRESHOLDING>
  <QUAD_MS_MS_TRANSMISSION>
    <LOW_MASS_ISOLATION_WINDOW VALUE=""/>
    <HIGH_MASS_ISOLATION_WINDOW VALUE=""/>
    <JOIN_ONLY_PCC_MATCHED_TO_SWITCHED_MASS VALUE="1"/>
    <SWITCHED_MASS_MATCH_TOL_PPM VALUE="100"/>
    <SWITCHED_MASS_CAL_OFFSET_PPM VALUE=""/>
  </QUAD_MS_MS_TRANSMISSION>
  <HYBRID_SONAR>
    <CALIBRATION_MODE VALUE="Auto"/>
    <MANUAL_CAL>
      <DRIFT_OFFSET VALUE="0"/>
      <DRIFT_SLOPE VALUE="0"/>
    </MANUAL_CAL>
  </HYBRID_SONAR>
```

```xml
<PRECURSOR_DEISOTOPING>
  <MIN_Z VALUE="2"/>
  <MAX_Z VALUE="6"/>
  <MIN_NUM_ISOTOPES VALUE=""/>
  <MZ_FWHM_WINDOW_FACTOR VALUE="0.5"/>
  <ISOTOPE_INTENSITY_RANGE_FACTOR VALUE="0.3333333"/>
  <MAKE_VIRTUAL_IONS VALUE="1"/>
  <DEISOTOPE_WITH_DRIFT>
    <INTENSITY_SAT_LOG10_RANGE VALUE="2"/>
    <MASS_DRIFT_TO_CHARGE_FRAC_MASS_TOL VALUE="50"/><!-- mDa -->
    <CHARGE_1_MAX_MZ_DRIFT_RATIO VALUE="8.8"/>
    <RING_DRIFT_FWHM_WINDOW_FACTOR VALUE="0.5"/>
    <RING_MZ_FWHM_WINDOW_FACTOR VALUE="1.333333"/>
    <RING_INTENSITY_HIGH_TIER_CEILING_FACTOR VALUE="0.50"/>
    <RING_INTENSITY_LOW_TIER_CEILING_FACTOR VALUE="0.15"/>
    <!-- Product Drift offset search range -->
    <PRODUCT_DRIFT_LOW_OFFSET VALUE="-4.5"/>
    <PRODUCT_DRIFT_HIGH_OFFSET VALUE="+0.5"/>
  </DEISOTOPE_WITH_DRIFT>
</PRECURSOR_DEISOTOPING>
<PRECURSOR_PEAK_TRACKING>
  <NO_PEAK_REJECTION_FOR_DDA_SPECTRA VALUE=""/>
  <ALIGN_MS1_INSOURCE_FRAGS_BY_PEAK_RET_TIME VALUE="0"/>
  <PROCESS_ONLY_TOP_N_INTENSE_PRECURSORS VALUE=""/>
  <EXTRACT_PRECURSORS_ABOVE_SNR VALUE="0">
    <SNR_THRESHOLD VALUE="0.59"/>
    <MIN_INTENSITY_CV_FOR_EXTRACTION VALUE="0.25"/>
```

```
  </EXTRACT_PRECURSORS_ABOVE_SNR>
  <PEAK_MIN_NUM_SCANS VALUE="5"/>
  <NUM_DROP_OUT_SCANS_TO_TERMINATE_PEAK_CHAIN VALUE="3"/>
  <MIN_PEAK_INTENSITY VALUE="16"/>
  <RATIO_OF_CHANGE_DECIMATION_FACTOR VALUE="2"/>
  <WRITE_PRECURSOR_PEAK_CHAINS_TO_CSV_FILE VALUE="1"/>
  <EXTRACT_PEAKS_WITHOUT_PEAK_PROPERTIES VALUE="1">
    <NUM_SCANS_FOR_PEAK_EXTRACTION_EACH_SIDE_OF_PEAK VALUE="3"/>
  </EXTRACT_PEAKS_WITHOUT_PEAK_PROPERTIES>
  <MIN_NUM_SCANS_FOR_LOW_HIGH_SMR_SELECTION VALUE="5"/>
  <SIGNAL_CV_THRESHOLD_FOR_LOW_HIGH_SMR_SELECTION VALUE="0.20"/>
  <LOW_CV_PEAK_SIGNAL_MEDIAN_RATIO_THRESHOLD VALUE="2.0"/>
  <HIGH_CV_PEAK_SIGNAL_MEDIAN_RATIO_THRESHOLD VALUE="1.0"/>
  <PEAK_MIN_NUM_SCANS_TO_REQUIRE_BOTH_HALF_HEIGHTS VALUE="21"/>
  <PEAK_BASE_WIDTH_EXTRACTION_IN_PLUS_MINUS_STD_DEVS VALUE="3.0"/>
  <ALLOW_PEAKS_WITH_NO_HALF_HEIGHTS VALUE="1"/>
  <MIN_PEAK_LEN VALUE="4"/>
  <MAX_PEAK_LEN VALUE="23"/>
</PRECURSOR_PEAK_TRACKING>
<ISOBARIC_LABEL_TRACKING>
  <TRACK_TMT_6PLEX_LABELS VALUE="0">
    <TMT_MASS_SEARCH_TOL_FACTOR VALUE="2.0"/>
  </TRACK_TMT_6PLEX_LABELS>
</ISOBARIC_LABEL_TRACKING>
<DRIFT_TRANSMISSION_ALIGNMENT VALUE="1">
  <!-- Determines coeluting precursors within drift (mass)
    transmission window for HDMSe/SONAR acquisitions
```

```
    and determines relative intensity ratio of each
    precursor then applies that ratio to their respective
    product ion spectrum as a ceiling threshold, extracting
    product ions within specified log intensity range. -->
  <DRIFT_TRANSMISSION_WINDOW_BINS VALUE="3"/>
  <PRODUCT_ION_TXM_INTENSITY_FACTOR VALUE="2.87"/>
  <PRODUCT_ION_LOG10_INTENSITY_RANGE VALUE="3"/>
</DRIFT_TRANSMISSION_ALIGNMENT>
<PRODUCT_ION_ALIGNMENT>
  <MIN_MZ VALUE="50"/>
  <REMOVE_RESIDUALS_FROM_PRODUCT_ION_SPECTRA VALUE="1"/>
  <REJECT_IONS_INTENSITY_GREATER_THAN_PRECURSOR VALUE="1"/>
  <FIX_SPLIT_IONS VALUE="1"/>
  <REMOVE_ISOTOPES VALUE="1"/>
  <CHARGE_REDUCE_PRODUCT_IONS_TO_MHPLUS VALUE="0"/>
  <ALIGN_PRODUCT_IONS_BY_INTENSITY_RATIO_OF_CHANGE VALUE="0">
    <PRECURSOR_TARGET_ROC VALUE="0.50"/>
    <PRECURSOR_PRODUCT_ION_ROC_MATCHING_FACTOR VALUE="0.40"/>
    <NUM_SCANS_BELOW_HALF_HEIGHTS_TO_STOP_ROC_COMPARES VALUE=""/>
    <TAKE_TOP_N_ROC_SCANS_CLOSEST_TO_TARGET VALUE=""/>
    <REJECT_PRODUCT_IONS_BY_ROC VALUE="0"/>
  </ALIGN_PRODUCT_IONS_BY_INTENSITY_RATIO_OF_CHANGE>
  <ALIGN_PRODUCT_IONS_BY_MASS_DEFECT VALUE="1">
    <B_ION_HIGH_STRIPE_FACTOR VALUE="1.25"/>
    <Y_ION_LOW_STRIPE_FACTOR VALUE="1.25"/>
  </ALIGN_PRODUCT_IONS_BY_MASS_DEFECT>
  <ALIGN_PRODUCT_IONS_BY_INTENSITY_MASS_BINS VALUE="0">
```

```
    <MASS_BIN_WIDTH_SET_BY_MR_DIVIDED_BY_NUM_BINS VALUE="20"/>
    <OVERRIDE_MASS_BIN_WIDTH_AMU VALUE="50"/>
    <KEEP_BIN_IF_NUM_IONS_LESS_THAN_EQ_TO VALUE="2"/>
    <REJECT_BIN_IF_INTENSITY_CV_LESS_THAN VALUE="0.25"/>
    <REJECT_ION_IF_INTENSITY_TO_MEAN_RATIO_LESS_THAN VALUE="0.50"/>
    <TAKE_TOP_N_INTENSE_BIN_IONS VALUE="5"/>
  </ALIGN_PRODUCT_IONS_BY_INTENSITY_MASS_BINS>
</PRODUCT_ION_ALIGNMENT>
<CPPIS_ALIGNMENT>
  <!-- MSe SSPPIS Construction Options: only one option can be set -->
  <MAKE_SSPPISS_FROM_MSE_SPECTRA_BETWEEN_HALF_HEIGHTS VALUE="0"/>
  <MAKE_SSPPISS_FROM_ALL_MSE_SPECTRA VALUE="1"/>
  <!-- SSPPIS Filter Options -->
  <USE_SSPPIS_IF_RESIDUAL_FRAG_EFF_GREATER_THAN VALUE=""/>
  <USE_SSPPIS_IF_NO_RESIDUAL_PCC_INTENSITY_GREATER_THAN VALUE=""/>
  <REJECT_SSPPIS_IF_PRECURSOR_INTENSITY_LESS_THAN_RESIDUAL VALUE="0"/>
  <COMPOSITING_MASS_MATCH_ERROR_TOL_PPM VALUE=""/>
  <MIN_NUMBER_OF_SPECTRA_FOR_CPPIS VALUE="3"/>
  <REJECT_COMPOSITE_ION_IF_AR1_CV_GREATER_THAN VALUE=""/>
  <CALC_COMPOSITE_PCC_ISOTOPE_RATIOS_AND_CV VALUE="1"/>
  <ALIGN_PRODUCT_IONS_BY_INTENSITY_MASS_BINS VALUE="1"/>
</CPPIS_ALIGNMENT>
<FILTERED_IONS_FUNC_CSV_FILES_PRODUCTION>
  <WRITE_FILTERED_IONS_FUNC_CSV_FILES VALUE="0"/>
  <WRITE_INCLUDE_EXCLUDE_IONS_AS_SEPARATE_FILES VALUE="1"/>
</FILTERED_IONS_FUNC_CSV_FILES_PRODUCTION>
<SSPPIS_MGF_PRODUCTION>
```

```xml
    <WRITE_SSPPISS_TO_MGF_FILE VALUE="1"/><!-- _iDTs_SSPPISs.mgf -->
    <WRITE_SSPPISS_ONLY_AT_PEAK_HEIGHT VALUE="0"/>
    <WRITE_PCC_ISOTOPE_RATIOS VALUE="1"/>
    <CHARGE_REDUCE_PEPMASS_TO_MHPLUS VALUE="0"/>
    <ADD_PRODUCT_ION_META_DATA VALUE="1"/>
    <SUPPRESS_VIRTUAL_PCC_SPECTRA VALUE="0"/>
    <SUPPRES_PRODUCT_IONS_WITH_ZERO_ROC VALUE="0"/>
    <MIN_NUMBER_PROD_IONS VALUE="3"/>
  </SSPPIS_MGF_PRODUCTION>
  <CPPIS_MGF_PRODUCTION>
    <WRITE_CPPISS_TO_MGF_FILE VALUE="1"/><!-- _iDTs_CPPISs.mgf -->
    <WRITE_PCC_ISOTOPE_RATIOS VALUE="1"/>
    <WRITE_PCC_ISOTOPE_RATIOS_CV VALUE="1"/>
    <CHARGE_REDUCE_PEPMASS_TO_MHPLUS VALUE="0"/>
    <ADD_PRODUCT_ION_META_DATA VALUE="1"/>
    <SUPPRESS_VIRTUAL_PCC_SPECTRA VALUE="0"/>
    <MIN_NUMBER_PROD_IONS VALUE="5"/>
  </CPPIS_MGF_PRODUCTION>
  <PLGS_PRODUCTION>
    <WRITE_CPPISS_TO_SPECTRUM_BIN_FILE VALUE="1"/>
    <COMBINE_CPPISS_INTO_AMRTS VALUE="0"/>
    <SPLIT_SPECTRUM_BIN_FILE_BY_CHARGE_STATES VALUE="0"/>
    <ADD_CHARGE1_SPECTRA_TO_SPLIT_SPECTRA VALUE="1"/>
    <SUPPRESS_VIRTUAL_PCC_SPECTRA VALUE="0"/>
    <MIN_NUMBER_CPPIS_PROD_IONS VALUE="5"/>
    <WRITE_LE_IDS_CPPIS_IDS_XREF_CSV_FILE VALUE="1"/>
  </PLGS_PRODUCTION>
```

```
  <PROGENESIS_PRODUCTION>
    <WRITE_CPPISS_TO_QI_FILES VALUE="0"/>
  </PROGENESIS_PRODUCTION>
</IDTS_RESEARCH_PARAMETERS>
endef

./ini/iDTs_control_2.xml :
    $$(file > ./ini/iDTs_control_2.xml,${iDTs_control_2})
```

The iDTs program is then executed for each file as follows:

```
define iDTs_rule_1
${1} &: ${2} ${3} ./ini/iDTs_control_1.xml
    cmd.exe /c ${idts_path} \
        -rawFile "$(call winify,$(2:.raw/_header.txt=.raw))" \
        -rawFileFuncCsvFiles "$(call winify,$(2:.raw/_header.txt=.raw))/Apex_1" \
        -paramsXmlFileName ./ini/iDTs_control_1.xml \
        -outputDir "$(call winify,$(2:.raw/_header.txt=.raw))/Apex_1/IDTs_1"
endef



$(foreach raw_iDTs, ${rawHead},$(eval $(call iDTs_rule_1,\
$(call idtsOut_1,${raw_iDTs}),\
${raw_iDTs},\
$(call apexOut_1,${raw_iDTs}))))
```

```
define iDTs_rule_2
${1} &: ${2} ${3} ./ini/iDTs_control_2.xml
    cmd.exe /c ${idts_path} \
                            -rawFile "$(call winify,$(2:.raw/_header.txt=.raw))" \
                            -rawFileFuncCsvFiles "$(call winify,$(2:.raw/_header.txt=.ra
                            -paramsXmlFileName ./ini/iDTs_control_1.xml \
                            -outputDir "$(call winify,$(2:.raw/_header.txt=.raw))/Apex_1
endef



$(foreach raw_iDTs, ${rawHead},$(eval $(call iDTs_rule_2,\
$(call idtsOut_2,${raw_iDTs}),\
${raw_iDTs},\
$(call apexOut_1,${raw_iDTs}))))
```

##### Search

The search is run through the Waters search system `iadb.exe`. The program requires an input of spectra from a Waters `.bin` file; a `.fasta` database to search; and a `.xml` file specifying the search parameters. The latter was exported from the PLGS gui and edited as below.

```
define searchSets
<?xml version="1.0" encoding="UTF-8"?>
<WORKFLOW_TEMPLATE TITLE="FBRH_BacteriaSearch" UUID="${UUID}"
            WORKFLOW_TEMPLATE_ID="_15731280752780_5292366780985168">
    <PROTEINLYNX_QUERY TYPE="Databank-search">
        <DATABANK_SEARCH_QUERY_PARAMETERS>
```

```
            <SEARCH_ENGINE_TYPE VALUE="PLGS"/>
            <SEARCH_TYPE NAME="Electrospray-Shotgun"/>
            <IA_PARAMS>
                <FASTA_FORMAT VALUE="Long Description"/>
                <PRECURSOR_MHP_WINDOW_PPM VALUE="6"/>
                <PRODUCT_MHP_WINDOW_PPM VALUE="8"/>
                <NUM_BY_MATCH_FOR_PEPTIDE_MINIMUM VALUE="6"/>
                <NUM_PEPTIDE_FOR_PROTEIN_MINIMUM VALUE="1"/>
                <NUM_BY_MATCH_FOR_PROTEIN_MINIMUM VALUE="9"/>
                <PROTEIN_MASS_MAXIMUM_AMU VALUE="300000"/>
                <FALSE_POSITIVE_RATE VALUE="1"/>
                <AQ_PROTEIN_ACCESSION VALUE=""/>
                <AQ_PROTEIN_MOLES VALUE="-1"/>
                <MANUAL_RESPONSE_FACTOR VALUE="-1"/>
                <DIGESTS>
                    <ANALYSIS_DIGESTOR MISSED_CLEAVAGES="1">
                        <AMINO_ACID_SEQUENCE_DIGESTOR NAME="Trypsin"
                                                    UUID="138fcbbc-4399-4fdb-ae63-9310f5de0f81">
                            <CLEAVES_AT AMINO_ACID="K" POSITION="C-TERM">
                                <EXCLUDES AMINO_ACID="P" POSITION="N-TERM"/>
                            </CLEAVES_AT>
                            <CLEAVES_AT AMINO_ACID="R" POSITION="C-TERM">
                                <EXCLUDES AMINO_ACID="P" POSITION="N-TERM"/>
                            </CLEAVES_AT>
                        </AMINO_ACID_SEQUENCE_DIGESTOR>
                    </ANALYSIS_DIGESTOR>
                </DIGESTS>
```

```
                <MODIFICATIONS>
                    <ANALYSIS_MODIFIER STATUS="FIXED">
                        <MODIFIER MCAT_REAGENT="No" NAME="Carbamidomethyl+C">
                            <MODIFIES APPLIES_TO="C" DELTA_MASS="57.0215"
                                                            TYPE="SIDECHAIN"/>
                        </MODIFIER>
                    </ANALYSIS_MODIFIER>
                    <ANALYSIS_MODIFIER ENRICHED="FALSE" STATUS="VARIABLE">
                        <MODIFIER MCAT_REAGENT="No" NAME="Oxidation+M">
                            <MODIFIES APPLIES_TO="M" DELTA_MASS="15.9949"
                                                        TYPE="SIDECHAIN"/>
                        </MODIFIER>
                    </ANALYSIS_MODIFIER>
                </MODIFICATIONS>
            </IA_PARAMS>
        </DATABANK_SEARCH_QUERY_PARAMETERS>
    </PROTEINLYNX_QUERY>
</WORKFLOW_TEMPLATE>
endef

./ini/iadb_params.xml : #.FORCE
    $$(file > ./ini/iadb_params.xml,${searchSets})
```

The search is then executed for each file with settings as below.

```
define iadb_rule
${1} &: ${2} ${3} ${combinedFasta} ./ini/iadb_params.xml
```

```
cmd.exe /c ${IADB_path} -RawFile "$(call winify,$(2:.raw/_header.txt=.raw))" \
    -pep3DFileName "$(call winify,$(filter %_iDTs_CPPIS_Spectrum.bin,${3}))" \
    -paraXMLFileName ./ini/iadb_params.xml \
    -proteinFASTAFileName ${combinedFasta} \
    -outPutDirName "$(dir $(call winify,$(filter %_iDTs_CPPIS_Spectrum.bin,${3})))"
    -outputUserDirName "$(dir $(call winify,$(filter %_iDTs_CPPIS_Spectrum.bin,${3})
    -pepMinAAcids 5 \
    -falsePositiveRatePass1 90 \
    -protFalsePositiveRateMax 50 \
    -precMuWindowPPM 40 \
    -prodMuWindowPPM 50 \
    -bMinBYNonRedundant 6 \
    -outputDistractionProteins 1 \
    -saveRaw 0 \
    -savePass0 0 \
    -savePass1 0 \
    -saveDebug 0 \
    -matchAllLossFrags 1 \
    -bDeveloperCSVOutput \
    -developer 1 \
    -allowHeMultiZ \
    -UW 1 \
    -WriteMGF 1 \
    -WriteXML 1 \
    -noPass2 0 \
    -newWorkflowXML 1 \
    -maxVarMods 2 \
```

```
        -bEnablePPMCalc 1 \
        -fragmentTypes BY \
        -bEnableFinalProtein 1 \
        -bEnableFinalPeptide 1 \
        -bEnableFinalFragment 1 \
        -bCollapseHomologs 0 \
        -saveSubsetDB 1

endef

localRaw:=${allRawData}Inj_40_Ecoli_F6_B4_7.raw/_header.txt



$(foreach raw_iadb,${rawHead},$(eval $(call iadb_rule,\
$(call iadbOut_1,${raw_iadb}),\
${raw_iadb},\
$(call idtsOut_1,${raw_iadb})\
)))



$(foreach raw_iadb,${rawHead},$(eval $(call iadb_rule,\
$(call iadbOut_2,${raw_iadb}),\
${raw_iadb},\
$(call idtsOut_2,${raw_iadb})\
)))
```

```
define mzidAllConvert
${2} : ${1}
    cmd.exe /c ${Csv2Mzid} -csv $(call winify,${1}) \
        -outputPath $(call winify,${2}) \
        -nologo -nomgf -pass2
endef

$(foreach file, ${iadb_all_final_frag},\
        $(eval $(call mzidAllConvert,\
        ${file},\
        $(file:.csv=.mzid))))
```

The resulting csv output file is the re-formatted to enable processing by the MAYU tool? from the trans-proteomics-pipeline using the R script as follows:

```
${Mayu_csv} : ${iadb_all_final_frag}
    ${R_path} --vanilla --quiet <<- EOF
```

```
    library('data.table')
    library('stringi')
    library('tidyverse')
    mayuAnnotateMods<-function(seq='MPCTEDYLSLILNR',
    mod='Carbamidomethyl+C(3);Oxidation+M(1)'){
    for(n in 1:length(seq)){
    #cat(n,'\n')
```

```
seqn<-seq[n]
modn<-mod[n]
if(modn!='None'){
modList<-stri_extract_all(modn,regex='[C|M]\\\\([\\\\d+|\\\\*]\\\\)',
simplify=TRUE)
modNum<-as.numeric(stri_extract(modList,regex='\\\\d+'))
modLt<-stri_extract(modList,regex='[C|M]')
replaceTab<-data.table(modNum,modLt)[order(-modNum)]
replaceTab[,modCode:=case_when(modLt=='C' ~ '=146.143',
                               modLt=='M' ~ '=147.192')]
print(replaceTab)
modSeq<-paste(replaceTab[order(modNum)][,code:=paste(modNum,modCode,sep='')][['code'
}else{
    modSeq<-''
}
if(!exists('outSeqVect')){outSeqVect<-modSeq}else{outSeqVect<-c(outSeqVect,modSeq)}
}
return(outSeqVect)
}
frac_list<-c(${iadb_all_final_frag_R})
fracAnnot<-paste(stri_extract(frac_list,regex='Inj_\\\\d+.*(?=.raw)'),
                        stri_extract(frac_list,regex='(?<=.raw\\\\/).*(?=\\\\/Inj)')
                        sep='_')
myFread<-function(input){fread(input,fill=TRUE)}
frac <- rbindlist(lapply(frac_list, myFread), idcol = 'rawFile', fill=TRUE)
frac[,DataSetLoc:=fracAnnot[rawFile]]
frac[,data:=stri_extract(DataSetLoc,regex='inj_\\\\d{2}+.+.(?=\\\\.raw)')]
```

```
frac[,opt:=stri_detect(DataSetLoc,fixed='_opt')]
fracAnnotMayu<-paste(stri_extract(frac_list,regex='Inj_\\\\d+.*(?=.raw)'),
                        stri_extract(frac_list,regex='(?<=.raw\\\\/).*(?=\\\\/Inj)')
                        sep='_')
frac_mayu <- rbindlist(lapply(frac_list, myFread), idcol = 'rawFile', fill=TRUE)
frac_mayu[,DataSetLoc:=fracAnnotMayu[rawFile]]
frac_mayu[,data:=stri_extract(DataSetLoc,regex='inj_\\\\d{2}+.+.(?=\\\\.raw)')]
frac_mayu[,opt:=stri_detect(DataSetLoc,fixed='_opt')]
write.csv(frac_mayu,file="frac_mayu.csv")
pepCheck_mu<-copy(frac_mayu)
myChooseHomolog<-function(x){
outSeqVect<-vector(length=0)
for(i in 1:length(x)){
cat('\n')
cat(x[i])
if(x[i]=='' | is.na(x[i]) ){
thisElement<-''
}else{
proteinList<-stri_split(x[i],fixed=' ',simplify=TRUE)
proteinList<-proteinList[!proteinList=='']
noRev<-proteinList[!stri_detect(proteinList, fixed='rev')]
if(length(noRev)!=0){
noCont<-noRev[!stri_detect(noRev, fixed='cont')]
if(length(noCont)!=0){
 thisElement<-sample(noCont, 1)
}else{
    #if noCont was zero length
```

```
    thisElement<-sample(noRev, 1)
  }
  }else{
  #if onRev was zero length
  thisElement<-sample(proteinList, 1)
  }
  }
  #cat(thisElement)
  outSeqVect<-c(outSeqVect,thisElement)
  }
  return(outSeqVect)
  }
  pepCheck_mu[,IDtype := case_when(stri_detect(pepCheck_mu[['protein.Accession']],fixe
              stri_detect(pepCheck_mu[['protein.Accession']],
              fixed='RANDOM') ~ 'RANDOM',
              stri_detect(pepCheck_mu[['protein.Accession']],
              fixed='halo') ~ 'Halo',
              stri_detect(pepCheck_mu[['protein.Accession']],
              regex='[OPQ][0-9][A-Z0-9]{3}[0-9]|[A-NR-Z][0-9]([A-Z][A-Z0-9]{2}[0-9]){1
              .default = 'Unknown')]
  pepCheck_mu[,pepUnkDat:=paste(rawFile,peptide.seq,precursor.z,precursor.mz,sep = '_'
  pepCheck_mu[,pepUnk:=paste(peptide.seq,precursor.z,precursor.mz,sep = '_')]
  pepCheck_mu[,mayuScanIndex:=paste('run',rawFile,'.',
                             round(((precursor.retT*60)/0.8)),
                             '.',
                             round(((precursor.retT*60)/0.8)),
                             '.',precursor.z,sep = '')][order(-peptide.score)]
```

```
pepCheck_mu[,cumHits:=1:dim(pepCheck_mu)[1]]
pepCheck_mu[,cumTarget:=cumsum(IDtype!='RANDOM')]
pepCheck_mu[,FDR:=100*(cumHits-cumTarget)/cumHits]
pepSum2<-pepCheck_mu[
    peptide.UniqueProducts>=5 | IDtype=="iRT"
    ][,.(peptide.seq=unique(.SD[which.max(peptide.score)][['peptide.seq']]),
    protein.Accession=unique(.SD[which.max(peptide.score)][['protein.Accession']]),
    protein.AccessionHomologue=unique(.SD[which.max(peptide.score)][['protein.Access
        pepMod=mayuAnnotateMods(unique(.SD[which.max(peptide.score)][['peptide.seq']
        unique(.SD[which.max(peptide.score)][['peptide.modification']])),
        peptide.score=unique(.SD[which.max(peptide.score)][['peptide.score']]),
        peptide.Rank=unique(.SD[which.max(peptide.score)][['peptide.Rank']]),
        peptide.UniqueProducts=unique(.SD[which.max(peptide.score)][['peptide.Unique
        by=mayuScanIndex][peptide.score>=4][order(-peptide.score)]
pepSum2[,cumHits:=1:dim(pepSum2)[1]]
pepSum2[,isDecoy:=stri_detect(protein.Accession, fixed='RANDOM')]
pepSum2[,cumTarget:=cumsum(!stri_detect(protein.Accession, fixed='RANDOM'))]
pepSum2[,FDR:=100*(cumHits-cumTarget)/cumHits]
pepSum2[,HomologShort:=myChooseHomolog(protein.AccessionHomologue)]
pepSum2[,protein.Accession:=case_when(protein.Accession!='' ~ protein.Accession,TRUE
write.table(pepSum2[!stri_detect(pepMod,fixed='NA'),
          c('mayuScanIndex','peptide.seq','protein.Accession','pepMod','peptide.sc
          file='${Mayu_csv}',
          row.names=FALSE, quote = FALSE, col.names=FALSE, sep=',')
EOF
```

MAYU requires the randomised protein squences identified by the search engine. Theses are extracted from the iadb results here and combined with true sequences to create a fasta file MAYU can used to estimate false discovery rates.

```
define seqkitFileList
./ini/seqkitFileList.txt :  ${subsetFasta}
    rm -f ./ini/seqkitFileList.txt
    $(foreach string,${subsetFasta},$(newline)$(tab)echo ${string} >> ./ini/seqkitFileLi
endef

$(eval $(call seqkitFileList,))


./ini/randomGrab.fasta : ./ini/seqkitFileList.txt ${subsetFasta}
    seqkit grep -o ./ini/randomGrab.fasta -r -p RANDOM --infile-list ./ini/seqkitFileLis
    sed -i 's/> R/>R/g' ./ini/randomGrab.fasta

${combinedFastaRand} : ${combinedFasta}
    shuffle -o ${combinedFastaRand} --nodesc ${combinedFasta}

${combinedFastaRandAnnotate} : ${combinedFastaRand}
    sed 's/>/>RANDOM_/g' ${combinedFastaRand} > ${combinedFastaRandAnnotate}

./ini/fastaForMayu.fasta : ${combinedFasta} ./ini/randomGrab.fasta ${combinedFastaRandAn
    cat ${combinedFasta} ./ini/randomGrab.fasta ${combinedFastaRandAnnotate} > ./ini/fas

./ini/fastaForMayu_noDup.fasta : ./ini/fastaForMayu.fasta
```

```
    seqkit rmdup -s -o ./ini/fastaForMayu_noDup.fasta -d ./ini/dupRecSeq.fasta ./ini/fas



${Mayu_out_txt} ${combinedMayu_psm} &: ${Mayu_csv} ./ini/fastaForMayu.fasta
    ${Mayu_path} -B ${Mayu_csv} -C "./ini/fastaForMayu.fasta" -M combinedMayu -H 1000 -E
```

The output from MAYU is processed using the following R script to create a csv file that can be used to build a spectral library for Skyline.

```
${final_frac_csv} ${final_frac_rds} &: ${iadb_all_final_frag} combinedMayu_psm_mFDR0.1_t
    ${R_path} --vanilla --quiet <<-EOF
```

```
    library('data.table')
    library('stringi')
    library('tidyverse')
    mayuMain<-fread('combinedMayu_main_1.07.csv')
    mayuPSM<-fread('combinedMayu_psm_mFDR0.1_t_1.07.csv')
    frac_list<-c(${iadb_all_final_frag_R})
    myFread<-function(input){fread(input,fill=TRUE)}
    fracAnnot<-stri_extract(frac_list,regex='Inj_\\\\d{1,2}.+\\\\.raw')
    frac <- rbindlist(lapply(frac_list, myFread), idcol = 'rawFile')
    frac[,DataSetLoc:=fracAnnot[rawFile]]
    frac[,data:=stri_extract(DataSetLoc,regex='inj_\\\\d{2}+.+.(?=\\\\.raw)')]
    frac[,pepUnkDat:=paste(rawFile,peptide.seq,precursor.z,precursor.mz,sep = '_')]
    frac[,pepUnk:=paste(peptide.seq,precursor.z,precursor.mz,sep = '_')]
    frac_lib<-frac[protein.Accession %in% c(mayuPSM[['prot']],'iRT')]
```

```
    pepList<-frac_lib[,.(
    pepUnkDat=.SD[which.max(peptide.score)][['pepUnkDat']],
    peptide.seq=.SD[which.max(peptide.score)][['peptide.seq']],
    peptide.score=.SD[which.max(peptide.score)][['peptide.score']]),
    by=pepUnk]
    candLib<-frac_lib[pepUnkDat %in% pepList[['pepUnkDat']] & peptide.matchType !="Adduc
    candLib<-candLib[ ,colnames(candLib) %in% names(frac), with=FALSE]
    write.csv(candLib,file='${final_frac_csv}',row.names = FALSE,quote = FALSE)
    saveRDS(frac,file='${final_frac_rds}')
    EOF
```

That library is then built using the build blib tool from Skyline.

```
${exp_blib} $(exp_blib:.blib=_redundant.blib) : ${final_frac_csv}
        cmd.exe /c ${buildblib_path} -P 0.001 -c 0 \
        $(call winify,${final_frac_csv}) \
        $(call winify,$(exp_blib:.blib=_redundant.blib))
        cmd.exe /c ${buildblibFilter_path} $(call winify,$(exp_blib:.blib=_redundant.bli
```

The library is converted to the trans proteomics pipeline splib format to facilitate wider re-use. The redundant `.blib` file is converted first to a `.ms2` and then to a `.splib` file. The `.splib` file is further processed to produce a consensus splib file.

```
${exp_splib} : $(exp_blib:.blib=_redundant.blib)
    cmd.exe /c ${blib2ms2_path} \
    -f $(call winify,$(exp_blib:.blib=.ms2)) $(call winify,$(exp_blib:.blib=_redundant.b
    ${spectrast_path} -cN$(exp_splib:.splib=_redundant) $(exp_blib:.blib=.ms2)
```

```
    ${spectrast_path} -cJU -cAC -cN$(exp_splib:.splib=_consensus) $(exp_splib:.splib=_re
    ${spectrast_path} -cAQ -cN$(exp_splib:.splib=) $(exp_splib:.splib=_consensus.splib)
```

##### Skyline Preprocessing

It turns out that skyline could not process the Waters "raw" format files if they included the mobility calibration files. If those files were excluded that data could be imported and mobility information was accurate. So for those files that were to be imported into Skyline coppies for the raw data excluding the mobility calibraion data files were created as follows:

```
define cpForSky
${1} : ${2}
    if [ ! -d "$(dir ${1})" ]; then mkdir -p $(dir ${1}); fi;
    cp -fT ${2} ${1}
endef

$(foreach file, ${rawFilesSkyline},$(eval $(call cpForSky,${file},$(call skyRawToOrigRaw
```

#### DIANN Processing

All data sets appart from the Vion were processed using DIA-NN v2.3.0. The setting for each data set are given in the comand below.

##### Create Library For Search

DIANN was used in library-free mode. So a single predicted library was generated for the fasta file, and that was used to search all data sets.

```
${combinedDIANNsplib} : ${combinedFasta}
    cmd.exe /c ${diann_path}  --lib "" \
            --threads 8 \
            --verbose 1 \
            --out-lib ${combinedDIANNsplib}
            --gen-spec-lib --predictor --reannotate \
            --fasta ${combinedFasta} \
            --fasta-search \
            --cut K*,R* \
            --min-fr-mz 200 --max-fr-mz 2000 \
            --min-pep-len 7 --max-pep-len 30 \
            --min-pr-mz 300 --max-pr-mz 2000 \
            --min-pr-charge 1 --max-pr-charge 4 \
            --cut K*,R* --missed-cleavages 1 \
            --var-mods 1 --unimod4 --var-mods 2 \
            -var-mod UniMod:35,15.994915,M \
            --var-mod UniMod:1,42.010565,*n \
            --met-excision
```

#### zenoTOF SWATH Data Processing with DIANN

##### Convert wiff to dia

For the Sciex data the “wiff” format was converted to “dia” format. This is not scrictly required since wiff data can be used directly by DIANN, but only if propriatry libraries are added to the DIANN folder. To make the data re-analysis univeraly available we convert to the “DIA” format here, which will be processable in all environments by DIANN.

```
define diannConvert
${1} : ${2}
    if [ ! -d "./dia" ]; then mkdir ./dia/; fi
    cmd.exe /c ${diann_path} --f $(call winify,$2) \
        --out-dir $(dir $(call winify,$1)) \
        --convert --threads 8 --verbose 1
endef

$(foreach file, ${wiff},\
        $(eval $(call diannConvert,\
        $(addprefix ./dia/,$(notdir $(file:.wiff=.wiff.dia))),\
        ${file})))
```

**zenoTOF Diann Processing**

```
# diann_zenoTOF wiff_dia

${diann_zenoTOF} &: ${wiff_dia}
    if [ ! -d "$(dir ${diannBaseRep_zeno})" ]; then mkdir -p $(dir ${diannBaseRep_zeno})
    if [ ! -d "$(dir ${diannBaseRep_zeno})tmp" ]; then mkdir -p $(dir ${diannBaseRep_zen
    cmd.exe /c ${diann_path} \
        $(foreach dia,${wiff_dia}, --f $(call winify,${dia})) \
        --lib ${combinedDIANNsplib} \
```

```
        --cont-quant-exclude cRAP- \
        --threads 8 --verbose 3 \
        --qvalue 0.01 --matrices \
        --reannotate \
        --fasta $(call winify,${combinedFasta}) \
        --min-fr-mz 100 --max-fr-mz 2000 --met-excision --cut K*,R* \
        --missed-cleavages 1 --min-pep-len 7 --max-pep-len 30 --min-pr-mz 400 \
        --var-mods 2 \
        --var-mod UniMod:35,15.994915,M --var-mod UniMod:1,42.010565,*n --unimod4 \
        --reanalyse --smart-profiling --pg-level 0 --export-quant \
        --temp $(dir ${diannBaseRep_timms})/tmp \
        --out  $(call winify,$(filter %-qn.parquet,${diann_zenoTOF})) \
        --out-lib $(call winify,$(filter %-lib.parquet,${diann_zenoTOF})) \
        --window 8 --mass-acc 20 --mass-acc-ms1 15 \
        --max-pr-mz 2000 --min-pr-charge 2 --max-pr-charge 4
```

#### Exploris DIA Processing with DIANN

##### Convert raw to dia

```
$(foreach file, ${ThermoRaw},\
        $(eval $(call diannConvert,\
        $(addprefix ./dia/,$(notdir $(file:.raw=.raw.dia))),\
        ${file})))
```

##### Exploris Diann Processing

```
${diann_ThermoRaw} &: ${ThermoRaw_dia}
    if [ ! -d "$(dir ${diannBaseRep_exploris})" ]; then mkdir -p $(dir ${diannBaseRep_ex
    if [ ! -d "$(dir ${diannBaseRep_exploris})tmp" ]; then mkdir -p $(dir ${diannBaseRep
    cmd.exe /c ${diann_path} \
        $(foreach dia,${ThermoRaw_dia}, --f $(call winify,${dia})) \
        --lib ${combinedDIANNsplib} \
        --cont-quant-exclude cRAP- \
        --threads 8 --verbose 3 \
        --qvalue 0.01 --matrices \
        --reannotate \
        --fasta $(call winify,${combinedFasta}) \
        --min-fr-mz 100 --max-fr-mz 2000 --met-excision --cut K*,R* \
        --missed-cleavages 1 --min-pep-len 7 --max-pep-len 30 --min-pr-mz 400 \
        --var-mods 2 \
        --var-mod UniMod:35,15.994915,M --var-mod UniMod:1,42.010565,*n --unimod4 \
        --reanalyse --smart-profiling --pg-level 0 --export-quant \
        --temp $(dir ${diannBaseRep_timms})/tmp \
        --out  $(call winify,$(filter %-qn.parquet,${diann_ThermoRaw})) \
        --out-lib $(call winify,$(filter %-lib.parquet,${diann_ThermoRaw})) \
        --window 8 --mass-acc 10 --mass-acc-ms1 10 \
        --max-pr-mz 2000 --min-pr-charge 2 --max-pr-charge 4
```

#### Timms DIA Processing with DIANN

##### Convert .d to dia

```
$(foreach file, ${Timms_d},\
        $(eval $(call diannConvert,\
        $(addprefix ./dia/,$(notdir $(file:.d=.d.dia))),\
        ${file})))
```

```
${diann_Timms_d} &: ${Timms_d_dia}
    if [ ! -d "$(dir ${diannBaseRep_timms})" ]; then mkdir -p $(dir ${diannBaseRep_timms
    if [ ! -d "$(dir ${diannBaseRep_timms})tmp" ]; then mkdir -p $(dir ${diannBaseRep_ti
    cmd.exe /c ${diann_path} \
        $(foreach dia,${Timms_d_dia}, --f $(call winify,${dia})) \
        --lib ${combinedDIANNsplib} \
        --cont-quant-exclude cRAP- \
        --threads 8 --verbose 3 \
        --qvalue 0.01 --matrices \
        --reannotate \
        --fasta $(call winify,${combinedFasta}) \
        --min-fr-mz 100 --max-fr-mz 2000 --met-excision --cut K*,R* \
        --missed-cleavages 1 --min-pep-len 7 --max-pep-len 30 --min-pr-mz 400 \
```

```
        --var-mods 2 \
        --var-mod UniMod:35,15.994915,M --var-mod UniMod:1,42.010565,*n --unimod4 \
        --reanalyse --smart-profiling --pg-level 0 --export-quant \
        --temp $(dir ${diannBaseRep_timms})/tmp \
        --out  $(call winify,$(filter %-qn.parquet,${diann_Timms_d})) \
        --out-lib $(call winify,$(filter %-lib.parquet,${diann_Timms_d})) \
        --window 8 --mass-acc 15 --mass-acc-ms1 15 \
        --max-pr-mz 2000 --min-pr-charge 2 --max-pr-charge 4
```

##### MRT

```
${diann_MRT} &: ${MRT_Raw}
    if [ ! -d "$(dir ${diannBaseRep_mrt})" ]; then mkdir -p $(dir ${diannBaseRep_mrt}); [illegible]
    if [ ! -d "$(dir ${diannBaseRep_mrt})tmp" ]; then mkdir -p $(dir ${diannBaseRep_mrt}[illegible]
    cmd.exe /c ${diann_path} \
        $(foreach mzml,${MRT_Raw}, --f $(call winify,${mzml})) \
        --lib ${combinedDIANNsplib} \
        --cont-quant-exclude cRAP- \
        --threads 8 --verbose 3 \
        --qvalue 0.01 --matrices \
        --reannotate \
        --fasta $(call winify,${combinedFasta}) \
        --min-fr-mz 100 --max-fr-mz 2000 --met-excision --cut K*,R* \
        --missed-cleavages 1 --min-pep-len 7 --max-pep-len 30 --min-pr-mz 400 \
        --var-mods 2 \
        --var-mod UniMod:35,15.994915,M --var-mod UniMod:1,42.010565,*n --unimod4 \
```

```
--reanalyse --smart-profiling --pg-level 0 --export-quant \
--temp $(dir ${diannBaseRep_mrt})/tmp \
--out  $(call winify,$(filter %-qn.parquet,${diann_MRT})) \
--out-lib $(call winify,$(filter %-lib.parquet,${diann_MRT})) \
--window 8 --mass-acc 5 --mass-acc-ms1 5 \
--max-pr-mz 2000 --min-pr-charge 2 --max-pr-charge 4
```

### Conclusion

The pipeline described above process data from the Waters Vion into the format where it is accessable to Skyline, and it processes all otheer data through DIANN and into the data format accessable to MSstats, which is described int he methods seciont in the main paper.
